# Cardiomyocyte-derived Wnt5a drives doxorubicin-induced cardiomyopathy by amplifying cellular senescence

**DOI:** 10.64898/2026.08.05.742723

**Authors:** Eun-Ah Sung, Peiyong Zhai, Soichiro Ikeda, Masato Matsushita, Koichiro Takayama, Takuma Takada, Nadezhda Fefelova, Lai-Hua Xie, Ghassan Yehia, Peter Romanienko, Varsha Gadiyar, Raymond Birge, Daniele Vecchio, Maurizio Forte, Valentina Valenti, Sebastiano Sciarretta, Junichi Sadoshima

**Affiliations:** Department of Cell Biology and Molecular Medicine, Cardiovascular Research Institute, Rutgers New Jersey Medical School, Newark, NJ, USA; Genome Editing Core Facility, Rutgers Cancer Institute of New Jersey, New Brunswick, NJ, USA; Department of Microbiology, Biochemistry and Molecular Genetics, Rutgers New Jersey Medical School, Newark, NJ, USA; Department of Medical and Surgical Sciences and Biotechnologies, Sapienza University of Rome, Latina, Italy; IRCCS Neuromed, Pozzilli, Italy; Department of Cardiology, ICOT University Hospital, Sapienza University of Rome, Latina, Italy

## Abstract

Doxorubicin (DOX) is an effective anthracycline chemotherapeutic agent, but its use is limited by cardiotoxicity that can progress to cardiomyopathy and heart failure. Cellular senescence contributes to DOX-induced cardiac injury, yet the upstream signals that initiate and propagate senescence in the injured heart remain unclear. Here, we identify Wnt5a, a non-canonical Wnt ligand, as a mediator of anthracycline cardiomyopathy. WNT5A was increased in serum from cancer patients receiving anthracycline therapy and in a pathologic human cardiomyocyte population in the context of DOX-induced cardiomyopathy. In mouse hearts, DOX induced early cardiomyocyte-enriched Wnt5a expression before overt cardiac dysfunction. Cardiomyocyte-specific *Wnt5a* deletion attenuated DOX-induced cardiac dysfunction, fibrosis and senescence marker induction, whereas recombinant Wnt5a and cardiomyocyte-targeted Wnt5a overexpression were sufficient to promote cardiomyocyte senescence and cardiac dysfunction. Mechanistically, DOX activated a Wnt5a-Fzd2 feed-forward axis that amplified Wnt5a expression in cardiomyocytes and propagated senescence to neighboring fibroblasts. Genetic disruption of this pathway in cardiomyocytes, fibroblasts or senescent cells reduced DOX-induced cardiomyopathy. Pharmacological inhibition of Wnt5a signaling with secreted frizzled-related protein 5 suppressed DOX-induced cardiac injury without compromising the anticancer efficacy of DOX. These findings identify Wnt5a-Fzd2 signaling as a senescence-amplifying mechanism in anthracycline cardiomyopathy and suggest a therapeutic strategy to mitigate DOX cardiotoxicity.

## Introduction

Anthracycline-based chemotherapy, particularly doxorubicin (DOX), remains a cornerstone of treatment for a broad range of malignancies, including lymphoma, leukemia, sarcoma, and breast cancer (1). However, its clinical use is limited by dose-dependent cardiotoxicity that can progress to cardiomyopathy and heart failure (1, 2). Patients who develop DOX-induced heart failure have poor outcomes, with a 5-year survival rate of approximately 50% (3, 4). Although DOX cardiotoxicity has been attributed to reactive oxygen species (ROS) generation and acute cardiomyocyte death (5, 6), antioxidant-based strategies have shown limited clinical benefit (7). Moreover, dexrazoxane, the only FDA-approved cardioprotective agent for anthracycline therapy, provides only partial protection and has raised concerns regarding potential interference with antitumor efficacy (8, 9). These limitations raise the need to define the mechanisms that drive the progressive and maladaptive cardiac remodeling underlying DOX cardiotoxicity.

Cellular senescence has emerged as an important contributor to DOX-induced cardiac injury (10–12). Senescence is characterized by persistent cell cycle arrest and acquisition of a proinflammatory senescence-associated secretory phenotype (SASP) (13). Although classically studied in proliferating cells, senescence can also be induced in postmitotic cardiomyocytes by genotoxic stress, including DOX exposure (10, 13–17). Importantly, pharmacological clearance of senescent cells improves cardiac function following DOX treatment even without replenishment of the eliminated cells, indicating that senescent cardiomyocytes contribute to cardiac dysfunction (10, 11, 18–20). These observations underscore the importance of identifying the upstream molecular pathways that induce the senescent phenotype in cardiomyocytes during DOX cardiotoxicity.

Wnt5a, a non-canonical Wnt ligand required during cardiac development, is expressed at low levels in the adult heart under basal conditions but is reactivated in response to pathological stress (21, 22). Increased Wnt5a expression has been reported in the myocardium and circulation of patients with heart failure and has been linked to adverse remodeling, including hypertrophy and fibrosis (23–26). Notably, recent transcriptomic analyses of human induced pluripotent stem cell (hiPSC)–derived cardiomyocytes from patients with DOX-induced heart failure identified elevated *WNT5A* expression (27). Despite these observations, whether Wnt5a directly contributes to DOX-induced cardiomyopathy and whether it promotes cardiomyocyte senescence remain unknown.

Here, we tested the hypothesis that DOX induces Wnt5a in the heart and that Wnt5a drives cardiac dysfunction by promoting cardiomyocyte senescence and the SASP. By defining the Wnt5a-senescence axis, this study identifies a mechanistic pathway underlying chemotherapy-associated cardiomyopathy and suggests a therapeutic framework for limiting cardiotoxicity while preserving the antitumor efficacy of DOX.

## Results

### Wnt5a is increased in anthracycline-induced cardiomyopathy

To determine whether Wnt5a, a highly conserved secreted protein, is associated with anthracycline-induced cardiomyopathy in humans, we measured circulating WNT5A levels in breast cancer patients receiving anthracycline therapy. Peripheral blood samples were collected from the same patients before treatment and after 3 months of anthracycline therapy (**Supplementary Table S1**). Serum WNT5A levels were significantly increased after anthracycline therapy compared to pretreatment baseline levels (**Figure 1A**).

**Figure 1.**
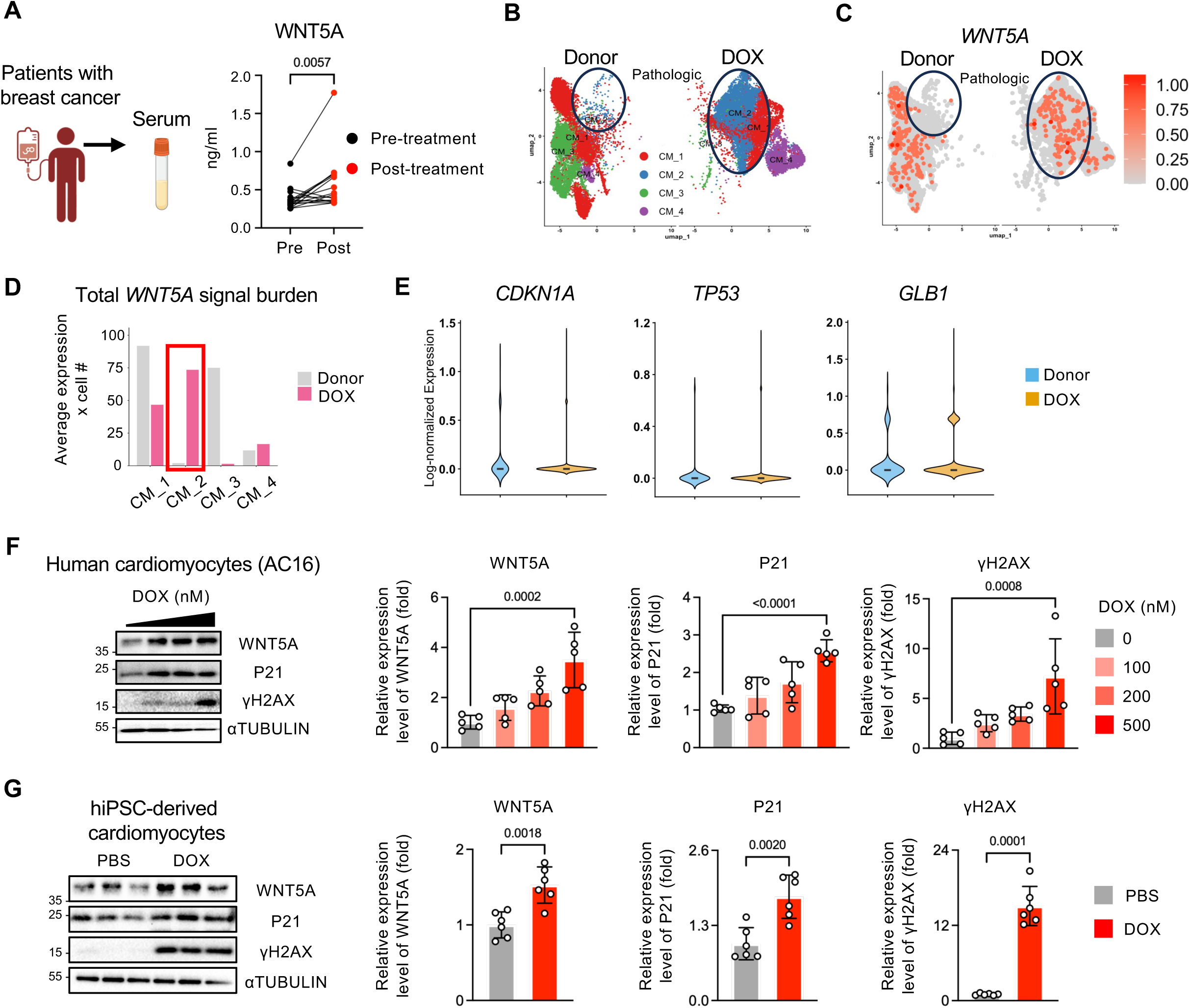
WNT5A is associated with human anthracycline cardiotoxicity and DOX-induced cardiomyocyte senescence. (**A**) Serum WNT5A levels in patients with breast cancer before and after anthracycline treatment, measured by ELISA. Paired samples are connected by lines. (**B**-**E**) Analysis of published snRNA-seq dataset from donor and DOX-induced cardiomyopathy hearts (GSE292067). UMAP plots of cardiomyocyte subclusters from donor and DOX-induced cardiomyopathy hearts in published human snRNA-seq dataset, showing cardiomyocyte subclusters (**B**) and *WNT5A* expression (**C**). (**D**) Total *WNT5A* signal burden across cardiomyocyte subclusters, calculated as average *WNT5A* expression x number of *WNT5A*-expressing cells. (**E**) Violin plots showing expression of *CDKN1A*, *TP53* and *GLB1* in CM_2 cardiomyocyte population from donor and DOX-induced cardiomyopathy hearts. (**F**) Human AC16 cardiomyocytes were treated with the indicated concentration of DOX for 48 hours. Cell lysates were subjected to Western blotting with anti-Wnt5a, p21, γH2AX and αTubulin antibodies. Relative expression levels of proteins were normalized by αTubulin. n = 5. (**G**) hiPSC-derived cardiomyocytes were treated with PBS or DOX (500 nM) for 48 hours. Cell lysates were subjected to Western blotting with anti-Wnt5a, p21, γH2AX and αTubulin antibodies. Relative expression levels of proteins were normalized by αTubulin. n = 6. Data are presented as mean ± SD. Statistical significance was determined by the Wilcoxon matched-paired signed-rank test (**A**), one-way ANOVA followed by multiple-comparison testing (**F**) and unpaired Student’s t-test (**G**). Exact P values are shown in the graphs.

To examine whether *WNT5A* is associated with a specific cardiomyocyte state in the DOX-treated human heart, we reanalyzed a recently published single-nucleus RNA sequencing (snRNA-seq) dataset from patients with DOX-induced cardiomyopathy and from nonfailing donor hearts (28). Cardiomyocytes were subclustered and annotated according to the previously reported marker-based classification, including *GRK5*-expressing CM_1, *NPPA*-expressing CM_2, *GRIK2*_expressing CM_3 and *TGFB2*-expressing CM_4 (**Supplementary Figure 1**) (28). Consistent with the original report, DOX-induced cardiomyopathic hearts showed expansion of the *NPPA*-expressing pathologic CM_2 population. (**Figure 1B**). Using this cardiomyocyte-state framework, we found that *WNT5A* signal was enriched in the expanded pathologic CM_2 population of DOX-treated patients but was comparatively limited in the CM_2 population of donor cardiomyocytes (**Figure 1C**). Quantification of the total *WNT5A* signal burden across cardiomyocyte states further showed that CM_2 is the major contributor to the *WNT5A* signal in DOX-induced cardiomyopathic hearts within the cardiomyocyte compartment (**Figure 1D**). We next assessed whether DOX CM_2 cells displayed a senescence-associated transcriptional profile. Compared with donor CM_2 cells, DOX CM_2 cells showed increased expression of *CDKN1A*, *TP53* and *GLB1* (**Figure 1E**).

Based on these human snRNA-seq findings, we next tested whether DOX directly induces WNT5A expression and senescence markers in human cardiomyocytes. AC16 human cardiomyocytes were treated with increasing concentrations of DOX. DOX increased WNT5A protein levels, together with senescence-associated markers, p21 and γH2AX, in AC16 human cardiomyocytes (**Figure 1F**). Consistently, DOX induced WNT5A protein expression and increased senescence-associated markers in human iPSC-derived cardiomyocytes (**Figure 1G**). Together, these findings show that WNT5A is increased in human anthracycline-induced cardiomyopathy and that DOX-induced WNT5A expression in human cardiomyocytes coincides with activation of senescence markers.

### DOX initially upregulates Wnt5a in the mouse heart in a cardiomyocyte-specific manner

Having established the clinical relevance of WNT5A in human anthracycline-induced cardiomyopathy, we investigated the temporal pattern and the cellular source of Wnt5a induction in an experimental mouse model of DOX-induced cardiomyopathy. C57BL/6J mice were treated with DOX using a well-established cardiomyopathy protocol consisting of weekly intraperitoneal injections of DOX at 5 mg/kg for 4 weeks, for a cumulative dose of 20 mg/kg (**Figure 2A**) (20). PBS-treated mice were used as controls. Wnt5a protein expression in mouse hearts was markedly increased as early as 7 days after DOX administration and remained elevated through day 28, the longest time point we examined (**Figure 2B**). Upregulation of senescence-associated markers became evident at later time points, reaching significance at week 2 for γH2AX and week 3 for p21. Both markers remained elevated at week 4 (**Figure 2B**). Echocardiographic analyses showed that left ventricular (LV) systolic dysfunction developed subsequently, with a significant reduction in LV ejection fraction by week 4 that persisted through week 12 (**Figure 2C**). These temporal changes suggest that Wnt5a induction is an early event that precedes the overt senescence marker induction and cardiac dysfunction after DOX treatment.

**Figure 2.**
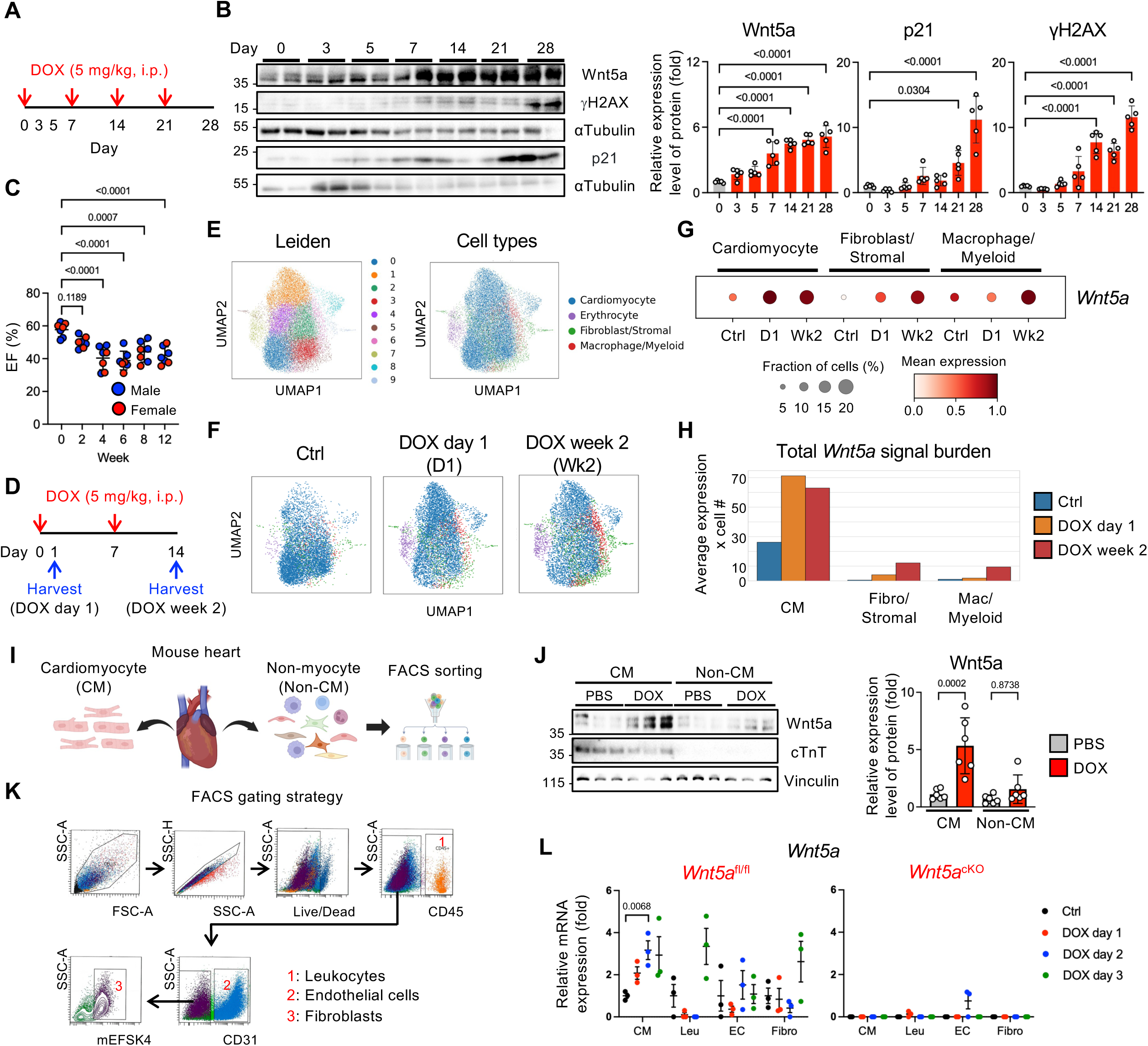
DOX initially upregulates Wnt5a predominantly in cardiomyocytes. (**A**-**C**) Wild-type mice were treated with DOX (5 mg/kg, i.p.) once weekly for 4 weeks. (**A**) Schematic of the DOX treatment protocol. (**B**) Heart lysates were subjected to Western blotting with anti-Wnt5a, p21, γH2AX and αTubulin antibodies at the indicated time points after DOX treatment. Protein levels were normalized by αTubulin. n = 5. (**C**) Left ventricular ejection fraction (LVEF) assessed by echocardiography. n = 7. (**D**-**H**) Mouse hearts were collected at day 1 after a single DOX injection or at week 2 after 2 weekly DOX injections and subjected to scRNA-seq analysis. (**D**) Experimental schematic. (**E**) UMAP plots showing Leiden clusters and major cardiac cell-type annotations from mouse heart scRNA-seq analysis. (**F**) UMAP plots showing major cardiac cell types. (**G**) Dot plot showing *Wnt5a* expression across major cardiac cell populations. Dot size indicates the fraction of *Wnt5a*-positive cells and color indicates mean Wnt5a expression. (**H**) Quantification of the total *Wnt5a* signal burden across major cardiac cell populations, calculated as average *Wnt5a* expression x number of *Wnt5a*-expressing cells. (**I**) Experimental schematic. (**J**) Cell lysates from isolated cardiomyocyte and non-myocyte fractions were subjected to Western blotting with anti-Wnt5a, cTnT and Vinculin antibodies. Protein levels were normalized by Vinculin. n = 6. (**K**) Representative FACS gating strategy for isolation of leukocytes, endothelial cells and fibroblasts from mouse hearts. (**L**) qPCR analysis of *Wnt5a* mRNA expression in sorted cardiac cell populations from *Wnt5a*^fl/fl^ and *Wnt5a*^cKO^ hearts after a single dose of DOX (5 mg/kg) treatment. n = 3. Data are presented as mean ± SD. Statistical significance was determined by one-way ANOVA followed by Tukey’s multiple-comparison test (**B, C, L**) and two-way ANOVA followed by Sidak’s multiple-comparison test (**J**). Exact P values are shown in the graphs.

To define the cellular source of Wnt5a during the early phase of DOX-induced cardiac injury, we performed single-cell RNA sequencing (scRNA-seq) of mouse hearts after DOX exposure. C57BL/6J mice received either a single dose of DOX (5 mg/kg) and hearts were collected on day 1, or two weekly doses of DOX and hearts were collected after 2 weeks (**Figure 2D**). After quality control and doublet removal, the data set was processed with batch correction and data integration using Harmony (29). Unsupervised clustering identified major cardiac cell populations, including cardiomyocytes, fibroblast/stromal cells, macrophage/myeloid cells and erythrocytes (**Figure 2, E and F and Supplemental Figure S2, A and B**). *Wnt5a* expression was preferentially increased in cardiomyocytes as early as day 1 after DOX treatment and remained detectable after 2 weeks (**Figure 2G**). Quantification of the total *Wnt5a* signal burden across major cardiac cell populations further showed that cardiomyocytes accounted for the majority of the *Wnt5a* signal following DOX treatment (**Figure 2H**). To validate this pattern at the biological replicate level, we calculated the percentage of *Wnt5a*-positive cardiomyocytes within each individual mouse. This mouse-level quantification showed an increased fraction of *Wnt5a*-positive cardiomyocytes after DOX treatment (**Supplementary Figure S2C**). These scRNA-seq findings suggest that DOX primarily induces early Wnt5a expression within the cardiomyocyte compartment.

To validate the cardiomyocyte origin of Wnt5a induction, we isolated cardiomyocyte and non-myocyte fractions from mouse hearts after DOX treatment, through enzyme digestion and separation with Percoll gradient (**Figure 2I**). Following a single DOX injection (5 mg/kg), Wnt5a protein expression was increased in cardiomyocytes but not in non-myocytes at day 5 (**Figure 2J**). We then generated cardiomyocyte-specific *Wnt5a* knockout mice (*Myh6*-*Cre Wnt5a*^fl/fl^, hereafter referred to as *Wnt5a*^cKO^ mice) to further test whether cardiomyocytes are the initial source of DOX-induced Wnt5a. To evaluate the level of Wnt5a in various cell types, mice were treated with a single dose of DOX (5 mg/kg) and cardiac cell populations were analyzed during the early injury phase (**Figure 2I**). Non-myocyte fractions were further sorted by FACS into leukocytes (CD45^+^), endothelial cells (CD45^-^CD31^+^) and fibroblasts (CD45^-^CD31^-^ mEFSK4^+^) (**Figure 2K**). qPCR analysis showed that *Wnt5a* mRNA was selectively increased in cardiomyocytes after DOX treatment, whereas leukocytes, endothelial cells and fibroblasts either did not show significant induction or exhibited delayed and lesser tendency of induction (**Figure 2L**). Importantly, DOX-induced *Wnt5a* upregulation was abolished across all cell fractions in *Wnt5a*^cKO^ mice (**Figure 2L**). Taken together, these findings identify cardiomyocytes as the early and predominant source of Wnt5a induction during DOX-induced cardiac injury.

### Wnt5a upregulation in cardiomyocytes is required for DOX-induced cardiomyopathy and senescence

To determine whether cardiomyocyte-derived Wnt5a is required for DOX-induced cardiomyopathy, *Wnt5a*^cKO^ mice and littermate *Wnt5a*^fl/fl^ controls were injected weekly with 5 mg/kg DOX for 4 weeks. Cardiac function was assessed by echocardiography 1 week after the final injection and hearts were harvested for biochemical and histological analyses. Western blot analyses showed that DOX-induced Wnt5a upregulation in the heart was nearly abolished in *Wnt5a*^cKO^ mice, further supporting cardiomyocytes as the primary source of DOX-induced cardiac Wnt5a expression (**Figure 3A**). The basal cardiac phenotype of *Wnt5a*^cKO^ mice did not differ from that of littermate *Wnt5a*^fl/fl^ mice in PBS-treated groups (**Figure 3B and Table 1**). Notably, however, DOX-induced LV dysfunction was markedly attenuated in *Wnt5a*^cKO^ mice compared to in *Wnt5a*^fl/fl^ mice (**Figure 3B**). Likewise, DOX-induced myocardial fibrosis was also significantly reduced in *Wnt5a*^cKO^ mice (**Figure 3C**).

**Figure 3.**
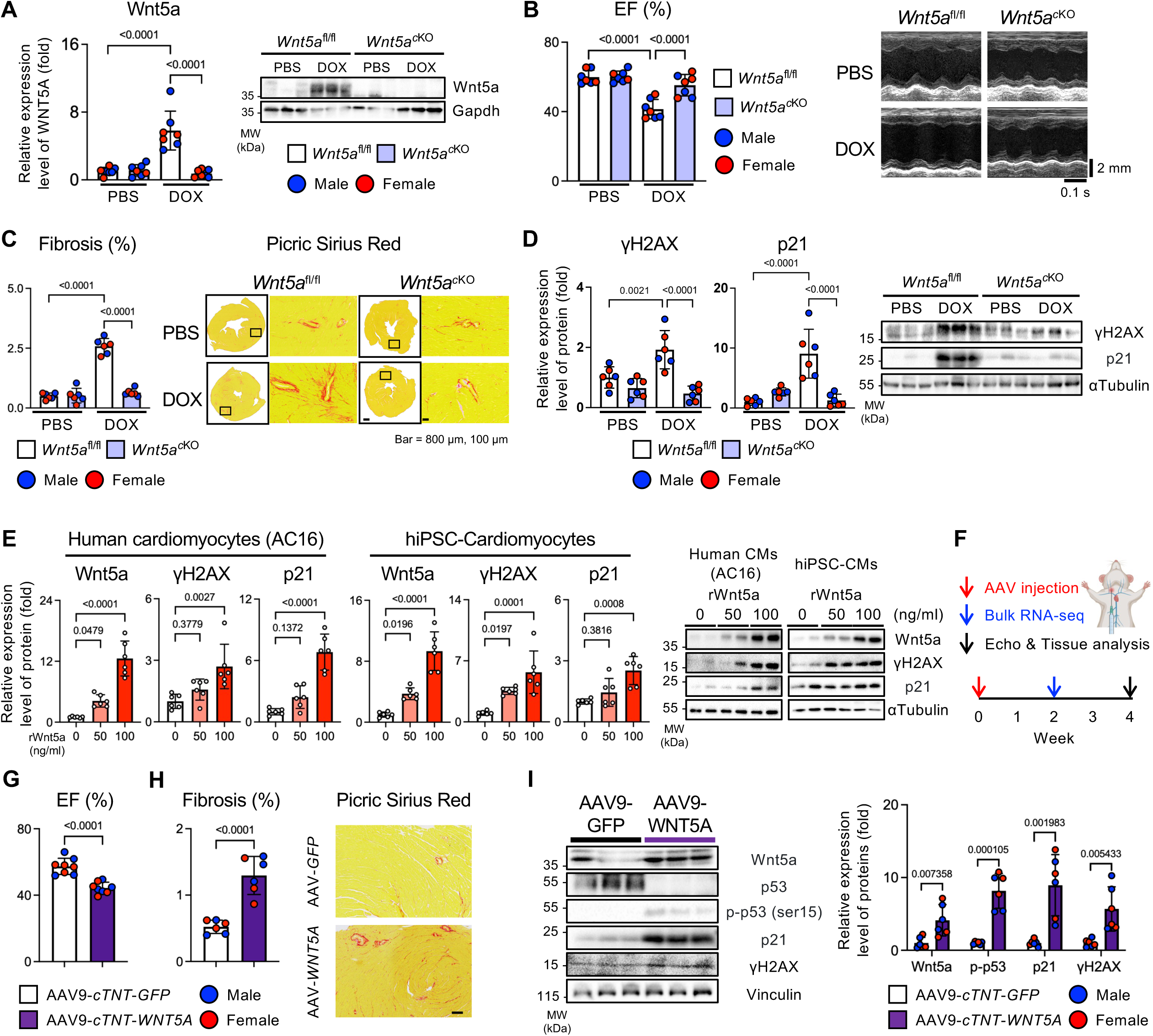
Cardiomyocyte-derived Wnt5a is necessary and sufficient for DOX-induced cardiac dysfunction and senescence. (**A**-**D**) *Wnt5a*^fl/fl^ and *Wnt5a*^cKO^ mice were treated with PBS or DOX (5 mg/kg, i.p.) once weekly for 4 weeks. One week after the final injection, echocardiography and heart tissue analysis were performed. (**A**) Heart lysates were subjected to Western blotting with anti-Wnt5a and Gapdh antibodies. Protein level was normalized by Gapdh. n = 6. (**B**) LVEF assessed by echocardiography and representative M-mode images are shown. n = 6. (**C**) Quantification and representative Picric Sirius Red staining of myocardial fibrosis from *Wnt5a*^fl/fl^ and *Wnt5a*^cKO^ mice. Scale bars: 800 μm (left) and 100 μm (right). n = 6. (**D**) Heart lysates were subjected to Western blotting with anti-γH2AX, p21 and αTubulin antibodies. Protein levels were normalized by αTubulin. n = 6. (**E**) AC16 cardiomyocytes and hiPSC-derived cardiomyocytes were treated with rWnt5a for 48 hours at the indicated concentrations. Cell lysates were subjected to Western blotting with anti-Wnt5a, γH2AX, p21 and αTubulin antibodies. Relative expression levels of proteins were normalized by αTubulin. n = 5. (**F**-**I**) Three-month-old C57BL/6J wild-type mice were subjected to intrajugular injection of AAV9-*cTNT*-*WNT5A* or AAV9-*cTNT*-*GFP*, followed by bulk RNA-seq, echocardiography and heart tissue analysis at the indicated time points. (**G**) LVEF assessed by echocardiography. n = 6. (**H**) Quantification and representative Picric Sirius Red-staining of myocardial fibrosis. Scale bar: 100 μm. n = 6. (**I**) Heart lysates were subjected to Western blotting with anti-Wnt5a, p53, p-p53 (Ser15), p21, γH2AX and Vinculin antibodies. Protein levels were normalized by Vinculin. n = 6. Data are presented as mean ± SD. Statistical significance was determined by two-way ANOVA followed by Sidak’s multiple-comparison test (**A**-**D**), one-way ANOVA followed by Tukey’s multiple-comparison test (**E**) and unpaired Student’s t-test (**G**-**I**). Exact P values are shown in the graphs.

**Table 1.** Echocardiographic measurements of *Myh6-*Cre/*Wnt5a*^fl/fl^ mice treated with DOX.

|  | *Wnt5a<sup>fl/fl</sup>* |  | *Myh6-Cre/Wnt5a<sup>fl/fl</sup>* |  |
| --- | --- | --- | --- | --- |
|  | PBS | DOX | PBS | DOX |
| n | 7 | 7 | 7 | 7 |
| IVSd (mm) | 0.58 ± 0.19 | 0.68 ± 0.08 | 0.61 ± 0.13 | 0.68 ± 0.17 |
| LVIDd (mm) | 3.71 ± 0.53 | 3.92 ± 0.19 | 3.58 ± 0.37 | 3.65 ± 0.50 |
| LVIDs (mm) | 2.55 ± 0.42 | 3.14 ± 0.26<sup>B</sup> | 2.45 ± 0.26 | 2.63 ± 0.42<sup>D</sup> |
| LVPWd (mm) | 0.65 ± 0.24 | 0.61 ± 0.05 | 0.75 ± 0.21 | 0.68 ± 0.10 |
| EF (%) | 59.90 ± 4.02 | 41.45 ± 5.61<sup>A</sup> | 60.18 ± 3.20 | 55.23 ± 6.20<sup>C</sup> |
| FS (%) | 31.31 ± 2.59 | 20.01 ± 3.12<sup>A</sup> | 31.40 ± 2.33 | 28.21 ± 3.86<sup>C</sup> |
| HR (bpm) | 437.02 ± 47.66 | 465.62 ± 45.60 | 438.71 ± 69.04 | 435.38 ± 37.89 |

Interventricular septal thickness (IVSd), Left ventricular internal diameter at end-diastole (LVIDd), Left ventricular internal diameter at end-systole (LVIDs), Left ventricular posterior wall thickness (LVPWd), Ejection fraction (EF), Fractional shortening (FS), Heart rate (HR). A, $P < 0.0001$ vs. *Wnt5a<sup>fl/fl</sup>*-PBS. B, $P < 0.01$ vs. *Wnt5a<sup>fl/fl</sup>*-PBS. C, $P < 0.001$ vs. *Wnt5a<sup>fl/fl</sup>*-DOX. D, $P < 0.05$ vs. *Wnt5a<sup>fl/fl</sup>*-DOX. Statistical significance was determined by unpaired Student's t-test. Data are presented as mean ± SD.

We next examined whether loss of cardiomyocyte-derived Wnt5a affects DOX-induced senescence-associated marker induction. DOX increased γH2AX and p21 protein levels in *Wnt5a*^fl/fl^ hearts, whereas these responses were blunted in *Wnt5a*^cKO^ hearts (**Figure 3D**). Consistently, *Wnt5a* knockdown in neonatal rat ventricular cardiomyocytes (NRVMs) reduced DOX-induced Wnt5a expression and attenuated induction of senescence markers (**Supplementary Figure 3A**). These results indicate that endogenous cardiomyocyte-derived Wnt5a is required for DOX-induced cardiac dysfunction, fibrosis, Wnt5a upregulation, and senescence.

### Wnt5a is sufficient to induce cardiomyocyte senescence and cardiac dysfunction

To investigate whether Wnt5a itself is sufficient to promote cardiomyocyte senescence, we treated AC16 human cardiomyocytes and hiPSC-derived cardiomyocytes with recombinant WNT5A (rWNT5A). rWNT5A treatment increased senescence markers, including p21 and γH2AX (**Figure 3E**). rWNT5A also increased WNT5A protein levels, suggesting that WNT5A amplifies its own expression in cardiomyocytes (**Figure 3E**). Similar effects were observed in isolated adult mouse cardiomyocytes, in which rWnt5a increased the number of γH2AX– and p21-positive cells (**Supplementary Figure 3B**). These results suggest that Wnt5a is sufficient to promote expression of senescence markers in human and mouse cardiomyocytes.

To determine whether cardiomyocyte-specific Wnt5a upregulation is sufficient to induce cardiac dysfunction *in vivo*, we injected AAV9-*cTNT*-*WNT5A* or AAV9-*cTNT*-*GFP* into the jugular veins of mice (**Figure 3F**). Increased Wnt5a expression (4.13-fold) in cardiomyocytes was sufficient to induce LV dysfunction compared to GFP-injected control mice (**Figure 3G**). Increased Wnt5a expression also elevated myocardial fibrosis (**Figure 3H**).

To gain insight into the transcriptional program activated by cardiomyocyte Wnt5a overexpression, we performed bulk RNA-sequencing (RNA-seq) of hearts from AAV9-*cTNT*-*WNT5A* and AAV9-*cTNT*-*GFP*-injected mice. Wnt5a overexpression increased expression of genes associated with the DNA damage response and the p53-p21 pathway, including *Atm*, *Rad9a*, *Rad9b*, *Parp1*, *Trp53*, *Pml* and *Cdkn1a*, while reducing expression of *Mdm4* (**Supplementary Figure S3C**). Consistent with these transcriptomic changes, AAV9-*cTNT*-*WNT5A*-injected mice showed increased protein levels of senescence markers, including p21 and γH2AX, and enhanced Ser15 phosphorylation of p53 (p-p53) in the heart (**Figure 3I**).

To explore a potential downstream signaling mechanism, we tested whether p38 MAPK contributes to Wnt5a-induced senescence-associated marker induction. Treatment with a p38 MAPK inhibitor reduced rWNT5A-induced increases in senescence markers, suggesting that p38 MAPK signaling may contribute to Wnt5a-mediated cardiomyocyte senescence (**Supplementary Figure S3D**).

Since senescent cells can influence neighboring cells through secretory factors, we further examined whether Wnt5a regulates SASP-related gene expression after DOX treatment. DOX increased the expression of multiple conventional SASP genes, including *Il1b*, *Il6*, *Il10*, *Tnfa* and *Ifng*, in *Wnt5a*^fl/fl^ control hearts, whereas these responses were attenuated in *Wnt5a*^cKO^ hearts (**Supplementary Figure S3E**). Likewise, DOX-induced expression of matrix-remodeling SASP genes, including *Mmp1a, Mmp1b* and *Mmp3,* was also reduced in *Wnt5a*^cKO^ hearts (**Supplementary Figure S3E**). We also assessed atypical SASP factors previously linked to aged cardiomyocytes, including *Gdf15*, *Edn3* and *Tgfb2* (17). Among these, only *Gdf15* was induced by DOX and reduced in *Wnt5a*^cKO^ hearts (**Supplementary Figure S3F**).

Collectively, these results demonstrate that cardiomyocyte-specific Wnt5a activation is sufficient to induce cardiac dysfunction, fibrosis, the DNA damage response and the SASP. Combined with the results obtained with *Wnt5a*^cKO^ mice, these results suggest that cardiomyocyte-derived Wnt5a is both necessary and sufficient for DOX-induced cardiomyopathy.

### Wnt5a production in p21^High^ senescent cells contributes to DOX-induced cardiomyopathy

Senescent cells can reinforce their own senescent state and promote senescence in neighboring cells through paracrine signaling (30, 31). Given that Wnt5a was induced early in cardiomyocytes and preceded the upregulation of senescence markers in response to DOX treatment, we next asked whether Wnt5a expression in senescent cells contributes to DOX-induced cardiomyopathy. To selectively target p21^High^ senescent cells *in vivo*, we used the *p21*^High^-Cre^ERT2^ mouse line, in which tamoxifen-inducible Cre recombinase is activated in a p21^High^ senescent cell-specific manner (32).

We first validated this system by crossing *p21*^High^-Cre^ERT2^ mice with *Rosa26-tdTomato* reporter mice. Flow cytometry analyses showed that, after 2 weeks of tamoxifen chow feeding followed by DOX treatment, tdTomato-positive cardiomyocytes were increased in DOX-treated hearts compared to in PBS-treated hearts (**Supplementary Figure S4A**). Moreover, tdTomato-positive cardiomyocytes exhibited γH2AX-positive nuclei, consistent with successful labeling of senescent cells by *p21*^High^-Cre^ERT2^ (**Figure 4A**). To further characterize these labeled cells, td-Tomato-negative and td-Tomato-positive cardiomyocytes were isolated from DOX-treated *p21*^High^-Cre^ERT2^/*Rosa26-tdTomato* mice by cell sorting. Senescence-related genes were significantly increased in td-Tomato-positive cells, compared to td-Tomato-negative cells (**Supplementary Figure S4B**). These findings indicate that tdTomato-positive cardiomyocytes represent a senescence-associated cardiomyocyte population after DOX injury.

**Figure 4.**
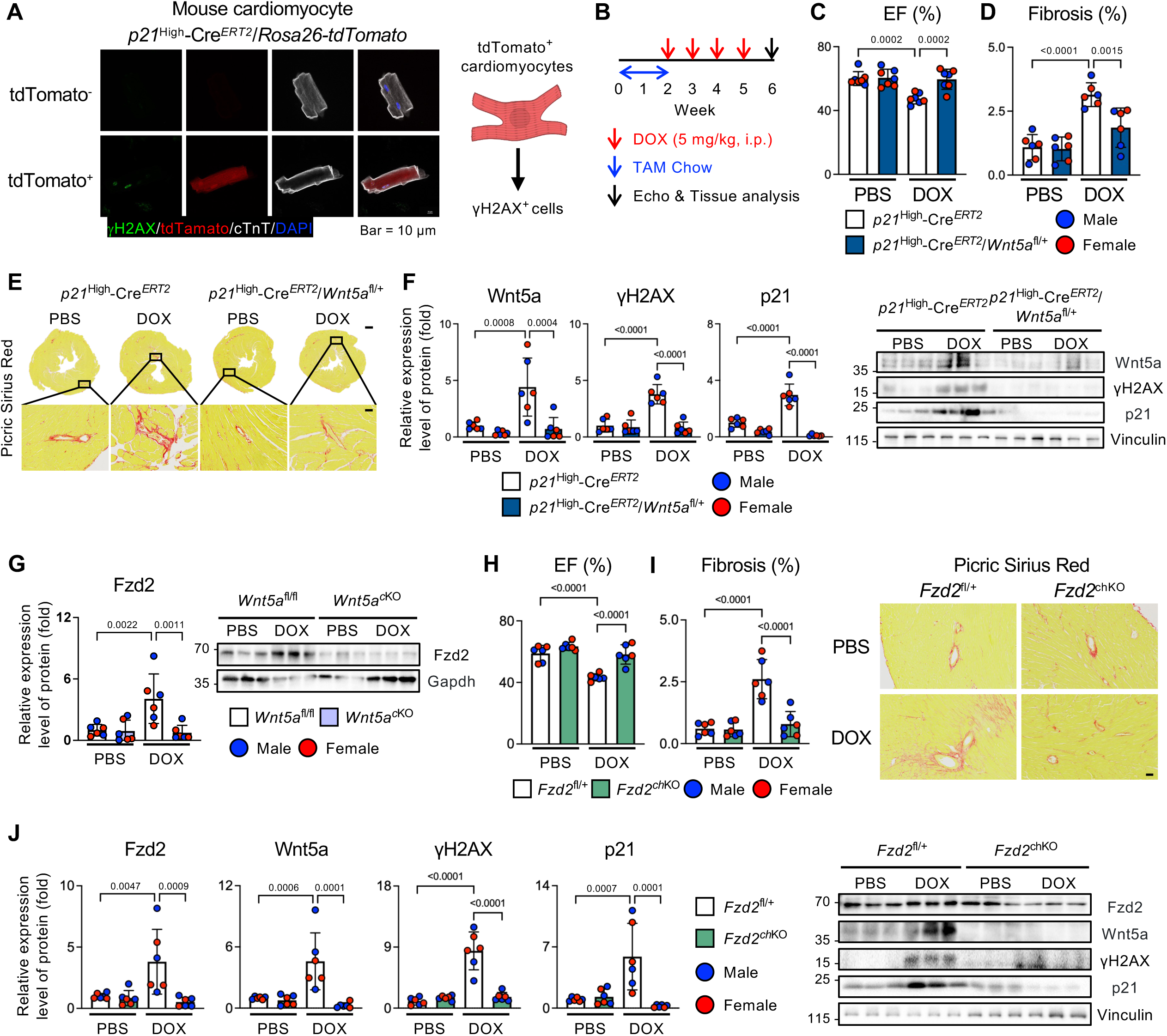
Wnt5a expression in *p21*^High^ cardiomyocytes and Fzd2 signaling contributes to DOX-induced cardiomyopathy. (**A**) Representative immunofluorescence images of tdTomato^-^ and tdTomato^+^ cardiomyocytes isolated from *p21*^High^-Cre^ERT2^;*Rosa26-tdTomato* mice. Cells were stained for γH2AX, cTnT and DAPI. Scale bar: 10 μm. (**B**-**F**) After two weeks of tamoxifen (TAM) chow feeding, *p21*^High^-Cre^ERT2^ and *p21*^High^-Cre^ERT2^/*Wnt5a*^fl/+^ mice were treated with PBS or DOX (5 mg/kg, i.p.) once weekly for 4 weeks. One week after the final injection, echocardiography and heart tissue analysis were performed. (**B**) Schematic of the experimental protocol. (**C**) LVEF was evaluated by echocardiography. n = 7. (**D**) Quantification of myocardial fibrosis. n = 7. (**E**) Representative Picric Sirius Red-stained heart sections. Scale bars: 500 μm (upper) and 50 μm (bottom). n = 6. (**F**) Heart lysates were subjected to Western blotting with anti-Wnt5a, γH2AX, p21 and Vinculin antibodies. Relative expression levels of proteins were normalized by Vinculin. n = 6. (**G**-**J**) *Fzd2*^fl/+^ mice and *Myh6*-Cre *Fzd2*^fl/+^ mice were treated with PBS or DOX (5 mg/kg, i.p.) once weekly for 4 weeks. One week after the final injection, echocardiography and heart tissue analysis were performed. (**G**) Heart lysates were subjected to Western blotting with anti-Fzd2 and Gapdh antibodies. Relative expression levels of Fzd2 were normalized to Gapdh. n = 6. (**H**) LVEF was evaluated by echocardiography. n = 6. (**I**) Quantification and representative Picric Sirius Red staining of myocardial fibrosis. Scale bar: 50 μm. n = 6. (**J**) Heart lysates were subjected to Western blotting with anti-Fzd2, Wnt5a, γH2AX, p21 and Vinculin antibodies. Relative expression levels of proteins were normalized by Vinculin. n = 6. Data are presented as mean ± SD. Statistical significance was determined by two-way ANOVA followed by Sidak’s multiple-comparison test. Exact P values are shown in the graphs.

To assess the role of Wnt5a in senescent cells, we crossed *p21*^High^-Cre^ERT2^ mice with *Wnt5a*^fl/+^ mice, enabling senescent cell-specific downregulation of *Wnt5a*. After 2 weeks of tamoxifen chow feeding, *p21*^High^-Cre^ERT2^/*Wnt5a*^fl/+^ mice and littermate *p21*^High^-Cre^ERT2^ controls (*p21*^High^-Cre^ERT2^ only) were treated with either DOX or PBS (**Figure 4B**). Downregulation of *Wnt5a* in p21^High^ cells significantly attenuated DOX-induced LV dysfunction and myocardial fibrosis (**Figure 4, C-E and Table 2**). In addition, DOX-induced increases in Wnt5a and senescent markers were also reduced in *p21*^High^-Cre^ERT2^/*Wnt5a*^fl/+^ hearts compared to in control hearts (**Figure 4F**).

**Table 2.** Echocardiographic measurements of *p21*^High^*-*Cre^ERT2^/*Wnt5a*^fl/+^ mice treated with DOX.

| | $p21^{\text{High-Cre}^{\text{ERT2}}}$ | | $p21^{\text{High-Cre}^{\text{ERT2}}/Wnt5a^{\text{fl/+}}}$ | |
| --- | --- | --- | --- | --- |
|  | PBS | DOX | PBS | DOX |
| n | 7 | 7 | 7 | 7 |
| IVSd (mm) | 0.60±0.11 | 0.64±0.05 | 0.58±0.05 | 0.62±0.08 |
| LVIDd (mm) | 4.05±0.32 | 4.42±0.26<sup>B</sup> | 3.90±0.34 | 3.99±0.29<sup>D</sup> |
| LVIDs (mm) | 2.76±0.21 | 3.37±0.19<sup>A</sup> | 2.66±0.32 | 2.74±0.34<sup>C</sup> |
| LVPWd (mm) | 0.58±0.11 | 0.61±0.07 | 0.58±0.08 | 0.60±0.05 |
| EF (%) | 60.03±4.31 | 47.44±3.31<sup>A</sup> | 60.50±5.40 | 59.59±6.30<sup>C</sup> |
| FS (%) | 31.67±3.20 | 23.69±2.04<sup>A</sup> | 31.93±3.73 | 31.39±4.14<sup>C</sup> |
| HR (bpm) | 445.70±80.80 | 433.75±22.02 | 447.46±63.37 | 435.20±80.25 |

Interventricular septal thickness (IVSd), Left ventricular internal diameter at end-diastole (LVIDd), Left ventricular internal diameter at end-systole (LVIDs), Left ventricular posterior wall thickness (LVPWd), Ejection fraction (EF), Fractional shortening (FS), Heart rate (HR). A, $P < 0.001$ vs. $p21^{\text{High-Cre}^{\text{ERT2}}}$ -PBS. B, $P < 0.05$ vs. $p21^{\text{High-Cre}^{\text{ERT2}}}$ -PBS. C, $P < 0.01$ vs. $p21^{\text{High-Cre}^{\text{ERT2}}}$ -DOX. D, $P < 0.05$ vs. $p21^{\text{High-Cre}^{\text{ERT2}}}$ -DOX. Statistical significance was determined by unpaired Student's t-test. Data are presented as mean ± SD.

scRNA-seq analysis showed that DOX increased the fraction of *Wnt5a^+^*cardiomyocytes in both the *Cdkn1a*^-^ and *Cdkn1a*^+^ populations, with *Cdkn1a* encoding the p21 protein (**Supplementary Figure S4, C and D**). *Wnt5a*^+^ cells were increased in *Cdkn1a*^-^ cardiomyocytes on day 1 after DOX treatment, indicating that Wnt5a induction is not restricted to *Cdkn1a*^+^ cardiomyocytes and may represent an early cardiomyocyte stress response. Notably, however, the fraction of *Wnt5a^+^* cells was greater in the *Cdkn1a*^+^ cardiomyocyte population than in the *Cdkn1a*^-^ cardiomyocyte population 2 weeks after DOX treatment. These findings suggest that Wnt5a is induced early after DOX exposure but becomes more enriched in the *Cdkn1a*^+^ senescent cardiomyocyte state during the later phase of injury.

Together, these results indicate that Wnt5a expression in p21^high^ cells contributes to DOX-induced cardiac dysfunction, fibrosis and senescence. These results also support a model in which Wnt5a participates in a feed-forward loop that reinforces senescence after DOX treatment.

### DOX induces Fzd2-dependent paracrine senescence signaling

We next investigated the identity of the Wnt receptor involved in DOX-induced Wnt5a signaling. Wnt5a, a ligand for both canonical and non-canonical Wnt pathways, binds to the Frizzled (Fzd) family and Ror receptors (33). Of the 10 Fzd isoforms and 2 Ror receptors examined, *Fzd2* was the only receptor significantly upregulated in mouse hearts after DOX treatment, as assessed by qPCR (**Supplementary Figure S4E**).

We further examined *Fzd2* expression in our mouse scRNA-seq dataset. Quantification of the total *Fzd2* signal burden across major cardiac cell populations showed that cardiomyocytes accounted for the majority of the *Fzd2* signal early after DOX treatment, beginning at day 1 (**Supplementary Figure S4F**). Consistent with this pattern, *Fzd2*^+^ cardiomyocytes were increased after DOX treatment (**Supplementary Figure S4G**). *Fzd2* signal also emerged in fibroblast/stromal cells at week 2, suggesting that the Wnt5a-Fzd2 signaling is initially upregulated in cardiomyocytes and later extends to non-myocyte populations during disease progression (**Supplementary Figure S4F**). Consistently, Fzd2 protein levels were increased in control hearts after DOX treatment, whereas this induction was blunted in *Wnt5a*^cKO^ hearts (**Figure 4F**). These results suggest that DOX coordinately induces both Wnt5a (ligand) and Fzd2 (receptor) in the heart and that Fzd2 upregulation is dependent on cardiomyocyte-derived Wnt5a.

To test whether DOX-treated cardiomyocytes exert paracrine effects, we performed conditioned medium transfer experiments. Conditioned medium was collected from NRVMs treated with DOX or PBS and then applied to separate recipient NRVMs (**Supplementary Figure S4H**). Conditioned medium from DOX-treated NRVMs increased Wnt5a and Fzd2 expression and induced senescence markers in recipient NRVMs (**Supplementary Figure S4H**). Silencing Fzd2 in recipient NRVMs attenuated these responses (**Supplementary Figure S4H**). These results suggest that soluble factors released from DOX-treated NRVMs promote senescence-associated signaling in recipient NRVMs through a Fzd2-dependent mechanism.

To elucidate the functional role of endogenous Fzd2 in mediating DOX-induced cardiotoxicity *in vivo*, we generated *Fzd2*^fl/+^ mice, which allow cell type-specific and conditional downregulation of Fzd2. By crossing *Fzd2*^fl/+^ mice with *Myh6*-*Cre* mice, we obtained cardiomyocyte-specific *Fzd2* heterozygous knockout (*Fzd2*^chKO^) mice. *Fzd2*^chKO^ mice and littermate *Fzd2*^fl/+^ controls were treated with DOX or PBS. There was no significant difference in ejection fraction between the PBS-treated groups, indicating preserved basal function in *Fzd2*^chKO^ mice (**Figure 4G and Table 3**). In contrast, the DOX-induced reduction in ejection fraction was markedly attenuated in *Fzd2*^chKO^ mice (**Figure 4H**). DOX-induced myocardial fibrosis was also reduced in *Fzd2*^chKO^ mice (**Figure 4I**). Moreover, the DOX-induced upregulation of Wnt5a, Fzd2 and senescence markers was decreased in *Fzd2*^chKO^ hearts (**Figure 4J**).

**Table 3.** Echocardiographic measurements of *Myh6-*Cre/*Fzd2*^fl/+^ mice treated with DOX.

|  | *Fzd2<sup>fl/+</sup>* |  | *Myh6-Cre/Fzd2<sup>fl/+</sup>* |  |
| --- | --- | --- | --- | --- |
|  | PBS | DOX | PBS | DOX |
| n | 6 | 6 | 6 | 6 |
| IVSd (mm) | 0.48±0.07 | 0.73±0.05<sup>A</sup> | 0.43±0.10 | 0.54±0.15<sup>D</sup> |
| LVIDd (mm) | 3.63±0.498 | 4.17±0.07<sup>B</sup> | 3.86±0.51 | 3.89±0.29<sup>D</sup> |
| LVIDs (mm) | 2.53±0.42 | 3.29±0.07<sup>B</sup> | 2.55±0.32 | 2.71±0.32<sup>C</sup> |
| LVPWd (mm) | 0.48±0.10 | 0.58±0.05<sup>B</sup> | 0.45±0.05 | 0.59±0.22 |
| EF (%) | 58.86±5.19 | 43.33±2.28<sup>A</sup> | 63.46±2.59 | 58.22±6.29<sup>C</sup> |
| FS (%) | 30.61±3.35 | 21.14±1.33<sup>A</sup> | 33.94±1.88 | 30.37±4.13<sup>C</sup> |
| HR (bpm) | 471.10±71.61 | 421.73±28.56 | 492.20±28.56 | 482.26±79.03 |

Interventricular septal thickness (IVSd), Left ventricular internal diameter at end-diastole (LVIDd), Left ventricular internal diameter at end-systole (LVIDs), Left ventricular posterior wall thickness (LVPWd), Ejection fraction (EF), Fractional shortening (FS), Heart rate (HR). A, P<0.0001 vs. *Fzd2<sup>fl/+</sup>*-PBS. B, P<0.05 vs. *Fzd2<sup>fl/+</sup>*-PBS. C, P<0.01 vs. *Fzd2<sup>fl/+</sup>*-DOX. D, P<0.05 vs. *Fzd2<sup>fl/+</sup>*-DOX. Statistical significance was determined by unpaired Student's t-test. Data are presented as mean ± SD.

Collectively, these results identify Fzd2 as a critical mediator of DOX-induced Wnt5a signaling in cardiomyocytes. The reduction of Wnt5a expression after Fzd2 downregulation further supports a feed-forward Wnt5a-Fzd2 signaling circuit that amplifies senescence and contributes to DOX-induced cardiomyopathy.

### Wnt5a propagates senescence to cardiac fibroblasts through Fzd2 and contributes to DOX-induced cardiomyopathy

Since Fzd2 signal emerged in fibroblast/stromal cells during the later phase of DOX injury, we asked whether the Wnt5a-Fzd2 signaling extends from cardiomyocytes to cardiac fibroblasts. Mouse scRNA-seq analyses showed that the fraction of *Fzd2*-positive fibroblast/stromal cells increased after DOX treatment, particularly at week 2 (**Supplementary Figure S5A**). To estimate potential Wnt5a-Fzd2 communication between cardiomyocytes and fibroblast/stromal cells, we calculated a mass-action based-ligand-receptor interaction score using average Wnt5a expression in sender cells and average Fzd2 expression in receiver cells. Based on the inferred communication map, Wnt5a-Fzd2 signaling was prominent as a cardiomyocyte autocrine loop at day 1 and fibroblast/stromal autocrine signaling became more evident by week 2, accompanied by increased cardiomyocyte-to-fibroblast/stromal communication (**Supplementary Figure S5, B and C**). These findings support a model in which DOX initially activates Wnt5a-Fzd2 signaling in cardiomyocytes and subsequently engages fibroblast/stromal Wnt5a-Fzd2 signaling during disease progression.

We next examined whether cardiomyocyte-derived Wnt5a exerts paracrine effects on cardiac fibroblasts *in vitro*. rWnt5a-treated neonatal rat cardiac fibroblasts (NRCFs) exhibited increased protein expression of Wnt5a and senescence markers (**Supplementary Figure S5D**). Similarly, conditioned medium from rWnt5a-treated NRVMs induced Wnt5a expression and senescence markers in NRCFs (**Supplementary Figure S5E**). In contrast, direct DOX treatment increased senescence markers but did not induce Wnt5a expression in fibroblasts, suggesting that Wnt5a upregulation in fibroblasts occurs preferentially through paracrine signaling from cardiomyocytes rather than through a direct fibroblast-autonomous response to DOX (**Supplementary Figure S5F**).

To investigate the functional relevance of endogenous fibroblast Fzd2 *in vivo*, we generated cardiac fibroblast-specific Fzd2-deficient mice by crossing *Fzd2*^fl/+^ mice with *Tcf21*-Cre^ERT2^ mice (hereafter referred to as *Fzd2*^cfhKO^ mice). After 2 weeks of tamoxifen chow feeding, *Fzd2*^cfhKO^ mice and littermate *Tcf21*-Cre^ERT2^ controls were treated with DOX or PBS (**Figure 5A**). Fibroblast-specific downregulation of Fzd2 significantly attenuated DOX-induced LV dysfunction and myocardial fibrosis (**Figure 5, B and C, Table 4**). These findings indicate that Fzd2 signaling in cardiac fibroblasts contributes to DOX-induced cardiac dysfunction.

**Figure 5.**
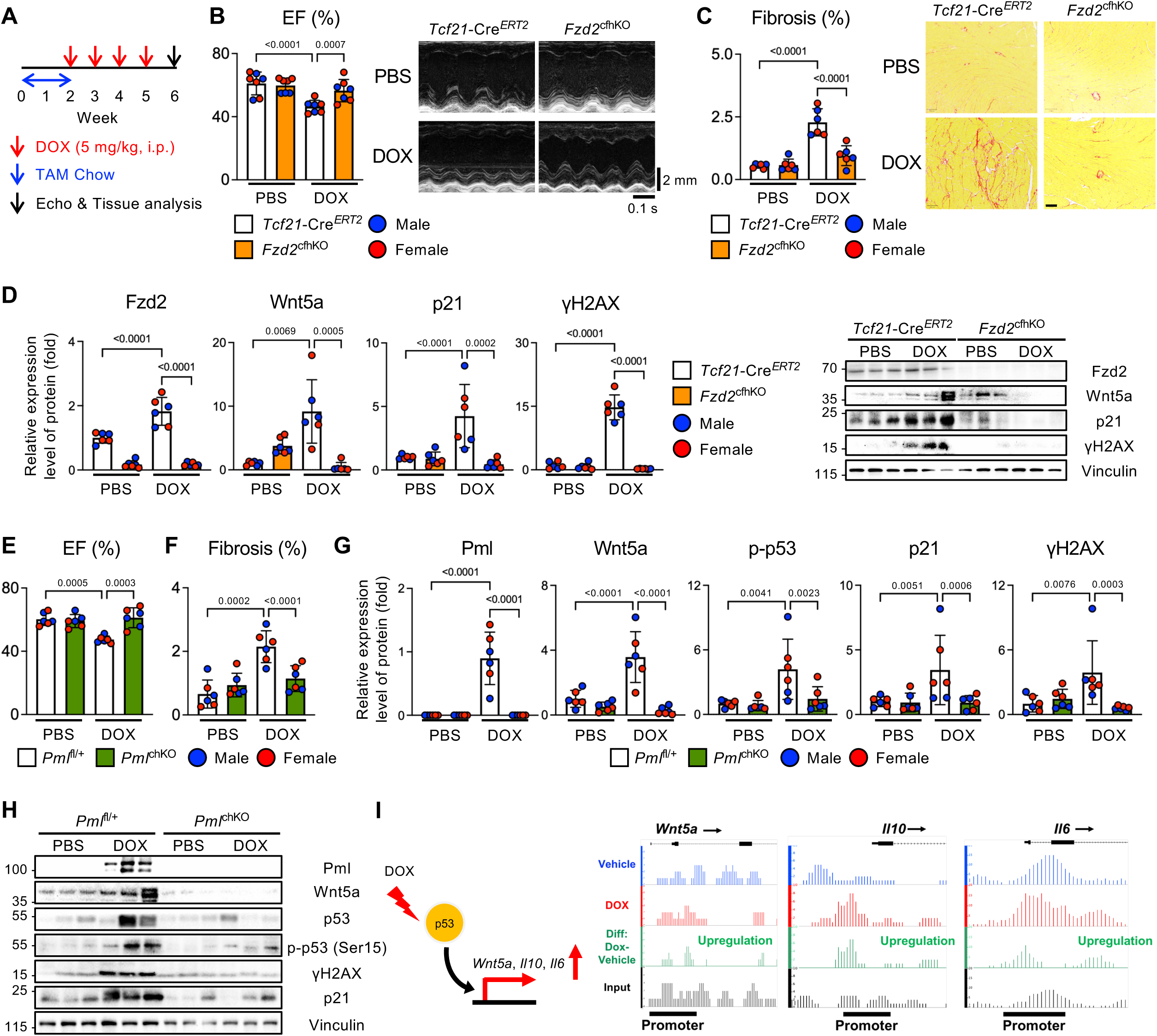
Fibroblast Fzd2 and cardiomyocyte Pml amplify Wnt5a-dependent senescence in DOX-induced cardiomyopathy. (**A**-**D**) After two weeks of tamoxifen (TAM) chow feeding, *Tcf21*-Cre^ERT2^ and *Tcf21*-Cre^ERT2^/*Fzd2*^fl/+^ mice were treated with PBS or DOX (5 mg/kg, i.p.) once weekly for 4 weeks. One week after the final injection, echocardiography and heart tissue analysis were performed. (**A**) Schematic of the experimental protocol. (**B**) LVEF was evaluated by echocardiography. n = 7. (**C**) Quantification of myocardial fibrosis and representative Picric Sirius Red-stained heart sections. Scale bar: 100 μm. n = 6. (**D**) Heart lysates were subjected to Western blotting with anti-Fzd2, Wnt5a, p21, γH2AX and Vinculin antibodies. Relative expression levels of proteins were normalized by Vinculin. n = 6. (**E**-**H**) *Pml*^fl/+^ mice and *Myh6*-Cre *Pml*^fl/+^ mice were treated with PBS or DOX (5 mg/kg, i.p.) once weekly for 4 weeks. One week after the final injection, echocardiography and heart tissue analysis were performed. (**E**) LVEF was evaluated by echocardiography. n = 6. (**F**) Quantification of myocardial fibrosis. n = 6. (**G** and **H**) Heart lysates were subjected to Western blotting with anti-Pml, Wnt5a, p53, p-p53 (Ser15), γH2AX, p21 and Vinculin antibodies. Relative expression levels of proteins were normalized by Vinculin. n = 6. (**I**) ChIP-seq analysis using an anti-p53 antibody in vehicle (PBS)– or DOX-treated mouse hearts (n = 4 per group). p53 ChIP-seq tracks showing p53 occupancy at the promoter regions of *Wnt5a*, *Il10* and *Il6* in vehicle– and DOX-treated hearts and green peaks representing differential enrichment relative to controls. Data are presented as mean ± SD. Statistical significance was determined by two-way ANOVA followed by Sidak’s multiple-comparison test. Exact P values are shown in the graphs.

**Table 4.** Echocardiographic measurements of *Tcf21*–Cre^ERT2^/*Fzd2*^fl/+^ mice treated with DOX.

|  | *Tcf21-Cre<sup>ERT2/+</sup>* |  | *Tcf21-Cre<sup>ERT2</sup>/Fzd2<sup>fl/+</sup>* |  |
| --- | --- | --- | --- | --- |
|  | PBS | DOX | PBS | DOX |
| n | 7 | 7 | 7 | 7 |
| IVSd (mm) | 0.50±0.08 | 0.65±0.09<sup>C</sup> | 0.47±0.05 | 0.49±0.03<sup>D</sup> |
| LVIDd (mm) | 3.57±0.58 | 3.96±0.34 | 3.98±0.24 | 3.57±0.69 |
| LVIDs (mm) | 2.41±0.37 | 3.05±0.24<sup>B</sup> | 2.73±0.16 | 2.56±0.61 |
| LVPWd (mm) | 0.50±0.11 | 0.56±0.07 | 0.45±0.03 | 0.48±0.07 |
| EF (%) | 61.06±7.23 | 46.35±4.00<sup>A</sup> | 59.79±4.57 | 56.49±7.07<sup>D</sup> |
| FS (%) | 32.18±5.10 | 22.83±2.46<sup>A</sup> | 31.44±3.23 | 28.98±4.29<sup>D</sup> |
| HR (bpm) | 417.22±50.87 | 444.85±29.87 | 408.52±48.91 | 424.37±80.36 |

Interventricular septal thickness (IVSd), Left ventricular internal diameter at end-diastole (LVIDd), Left ventricular internal diameter at end-systole (LVIDs), Left ventricular posterior wall thickness (LVPWd), Ejection fraction (EF), Fractional shortening (FS), Heart rate (HR). A, P<0.001 vs. *Tcf21-Cre<sup>ERT2</sup>*-PBS. B, P<0.01 vs. *Tcf21-Cre<sup>ERT2</sup>*-PBS. C, P<0.05 vs. *Tcf21-Cre<sup>ERT2</sup>*-DOX. D, P<0.01 vs. *Tcf21-Cre<sup>ERT2</sup>*-DOX. Statistical significance was determined by unpaired Student's t-test. Data are presented as mean ± SD.

Notably, *Fzd2*^cfhKO^ mice showed reduced DOX-induced expression of senescence markers, together with suppression of Fzd2 and Wnt5a protein induction (**Figure 5D**). The reduction in Wnt5a expression suggests that fibroblast Fzd2 contributes to secondary amplification of Wnt5a signaling in the DOX-treated heart. These results indicate that inhibition of senescence propagation via blockade of the Wnt5a receptor Fzd2 in fibroblasts is sufficient to mitigate DOX-induced cardiotoxicity, even when cardiomyocyte-derived Wnt5a remains intact. Taken together, these findings support a model in which cardiomyocyte-derived Wnt5a may engage Fzd2 on cardiac fibroblasts, promoting secondary fibroblast activation and the amplification of senescence during DOX-induced cardiac injury.

### A reciprocal Wnt5a-Pml-p53 circuit reinforces DOX-induced senescence in the heart

We next investigated the mechanism linking Wnt5a induction to p53-p21 activation during DOX-induced cardiac injury. As shown above, cardiomyocyte-targeted Wnt5a upregulation induced p53-p21 pathway genes and proteins *in vivo* (**Figure 3I and Supplementary Figure S3C)**. Consistent with activation of this pathway, scRNA-seq analysis of mouse hearts showed that DOX-treated *Cdkn1a*^+^ cardiomyocytes exhibited increased expression of selected p53 target and stress-response genes, including *Mdm2*, *Gadd45a*, *Sesn1* and *Trp53inp1* (**Supplementary Figure S5G**).

Promyelocytic leukemia protein (Pml) has been implicated in p53-dependent senescence (34, 35). Pml and p53 physically interact and mutually reinforce one another’s activity, thereby promoting senescence together (36). In our AAV9-*cTNT*-*WNT5A* bulk RNA-seq dataset, *Pml* was increased in mouse hearts with *Wnt5a* upregulation in cardiomyocytes, suggesting that Pml may be involved in Wnt5a signaling (**Supplementary Figure S3C**). To test whether Pml contributes to DOX-induced cardiomyocyte senescence, we knocked down *Pml* in NRVMs. DOX increased Pml protein expression, whereas *Pml* knockdown attenuated DOX-induced expression of p-p53 and p21 proteins (**Supplementary Figure S5H**). Notably, *Pml* knockdown also reduced DOX-induced Wnt5a protein expression, suggesting that Pml contributes not only to p53-p21 activation but also to the maintenance of Wnt5a expression during DOX-induced injury (**Supplementary Figure S5H**). These findings suggest that Pml is not simply a downstream effector of DOX-induced senescence signaling but rather functions within a reciprocal Wnt5a-Pml-p53 circuit that sustains Wnt5a expression and reinforces p53-p21 activation in NRVMs.

To determine the *in vivo* relevance of Pml in DOX-induced cardiotoxicity, we generated cardiomyocyte-specific *Pml* heterozygous knockout mice by crossing *Pml*^fl/+^ mice with *Myh6*-Cre mice (hereafter referred to as *Pml*^chKO^ mice). *Pml*^chKO^ mice showed preservation of LV systolic function compared with littermate controls, indicating protection from DOX-induced cardiac dysfunction (**Figure 5E and Table5**). DOX-induced myocardial fibrosis was also decreased in *Pml*^chKO^ hearts (**Figure 5F and Supplementary Figure S5I**). DOX increased Pml protein expression in control hearts, whereas this induction was markedly attenuated in *Pml*^chKO^ hearts (**Figure 5G**). DOX-induced increases in Wnt5a, p-p53, p-53, p21 and γH2AX proteins were also reduced in *Pml*^chKO^ hearts (**Figure 5, G and H**). Together, these data indicate that cardiomyocyte Pml contributes to DOX-induced Wnt5a induction, p53-p21 activation and cardiac dysfunction.

**Table 5.** Echocardiographic measurements of *Myh6*–Cre*Pm*/^fl/+^ mice treated with DOX.

|  | *Pml<sup>fl/+</sup>* |  | *Myh6-Cre/Pml<sup>fl/+</sup>* |  |
| --- | --- | --- | --- | --- |
|  | PBS | DOX | PBS | DOX |
| n | 6 | 6 | 6 | 6 |
| IVSd (mm) | 0.60±0.17 | 0.53±0.10 | 0.50±0.15 | 0.57±0.20 |
| LVIDd (mm) | 4.23±0.32 | 4.20±0.29 | 4.08±0.22 | 3.92±0.37 |
| LVIDs (mm) | 2.88±0.27 | 3.21±0.22<sup>B</sup> | 2.86±0.15 | 2.64±0.22<sup>C</sup> |
| LVPWd (mm) | 0.53±0.10 | 0.49±0.07 | 0.53±0.12 | 0.57±0.17 |
| EF (%) | 60.26±3.97 | 47.41±2.22<sup>A</sup> | 57.70±2.50 | 61.21±6.27<sup>C</sup> |
| FS (%) | 31.92±2.74 | 23.55±1.40<sup>A</sup> | 30.01±1.72 | 32.49±4.50<sup>C</sup> |
| HR (bpm) | 462.22±43.00 | 451.01±31.06 | 441.27±42.82 | 443.76±20.28 |

Interventricular septal thickness (IVSd), Left ventricular internal diameter at end-diastole (LVIDd), Left ventricular internal diameter at end-systole (LVIDs), Left ventricular posterior wall thickness (LVPWd), Ejection fraction (EF), Fractional shortening (FS), Heart rate (HR). A, P<0.0001 vs. *Pml<sup>fl/+</sup>*-PBS. B, P<0.05 vs. *Pml<sup>fl/+</sup>*-PBS. C, P<0.001 vs. *Pml<sup>fl/+</sup>*-DOX. Statistical significance was determined by unpaired Student's t-test. Data are presented as mean ± SD.

We asked whether p53 directly regulates transcriptional programs linked to Wnt5a and the senescence-associated secretory signaling in DOX-injured hearts. ChIP-seq analyses using an anti-p53 antibody were performed on hearts from PBS– and DOX– treated wild-type mice. DOX increased p53 occupancy at the promoter regions of several SASP genes, including *Wnt5a*, *Il6* and *Il10* (**Figure 5I**). These findings suggest that DOX enhances p53-dependent transcriptional regulation of *Wnt5a* and SASP-related genes, providing a mechanism by which Pml-p53 signaling reinforces Wnt5a expression and senescence paracrine signaling during DOX injury.

Together, these results identify Pml as a key component of the Wnt5a-Pml-p53 feedback circuit that sustains Wnt5a expression, reinforces p53-p21 activation and promotes DOX-induced cardiomyopathy.

### sFRP5 suppresses DOX-induced cardiomyocyte senescence and protects against cardiotoxicity without compromising anti-tumor efficacy

Secreted frizzled-related protein 5 (sFRP5) is an endogenous Wnt antagonist that can inhibit Wnt5a signaling by preventing ligand engagement with Fzd receptors (37). We tested whether pharmacological inhibition of the Wnt5a-Fzd2 pathway could suppress DOX-induced senescence. Human AC16 cardiomyocytes and hiPSC-derived cardiomyocytes were treated with recombinant sFRP5 in the presence or absence of DOX. sFRP5 inhibited DOX-induced increases in γH2AX and p21 protein levels (**Figure 6, A and B**). Similarly, sFRP5 also inhibited DOX-induced increases in γH2AX and p21 protein levels in NRVMs (**Supplementary Figure 6A**). Moreover, adult mouse cardiomyocytes isolated from *p21*^High^-Cre^ERT2^;*Rosa26-tdTomato* mice showed a reduction in DOX-induced tdTomato-positive p21^High^ cardiomyocytes after sFRP5 treatment (**Figure 6C**). These findings indicate that sFRP5 suppresses DOX-induced senescence marker induction in cardiomyocytes in both humans and rodents. To assess whether systemic sFRP5 administration reduces DOX-induced senescence *in vivo*, wild-type mice were treated with DOX in the presence or absence of recombinant sFRP5 (**Figure 6D**). sFRP5 attenuated DOX-induced increases in Wnt5a and senescence marker expression, including p21 and γH2AX, in the mouse hearts (**Figure 6D**).

**Figure 6.**
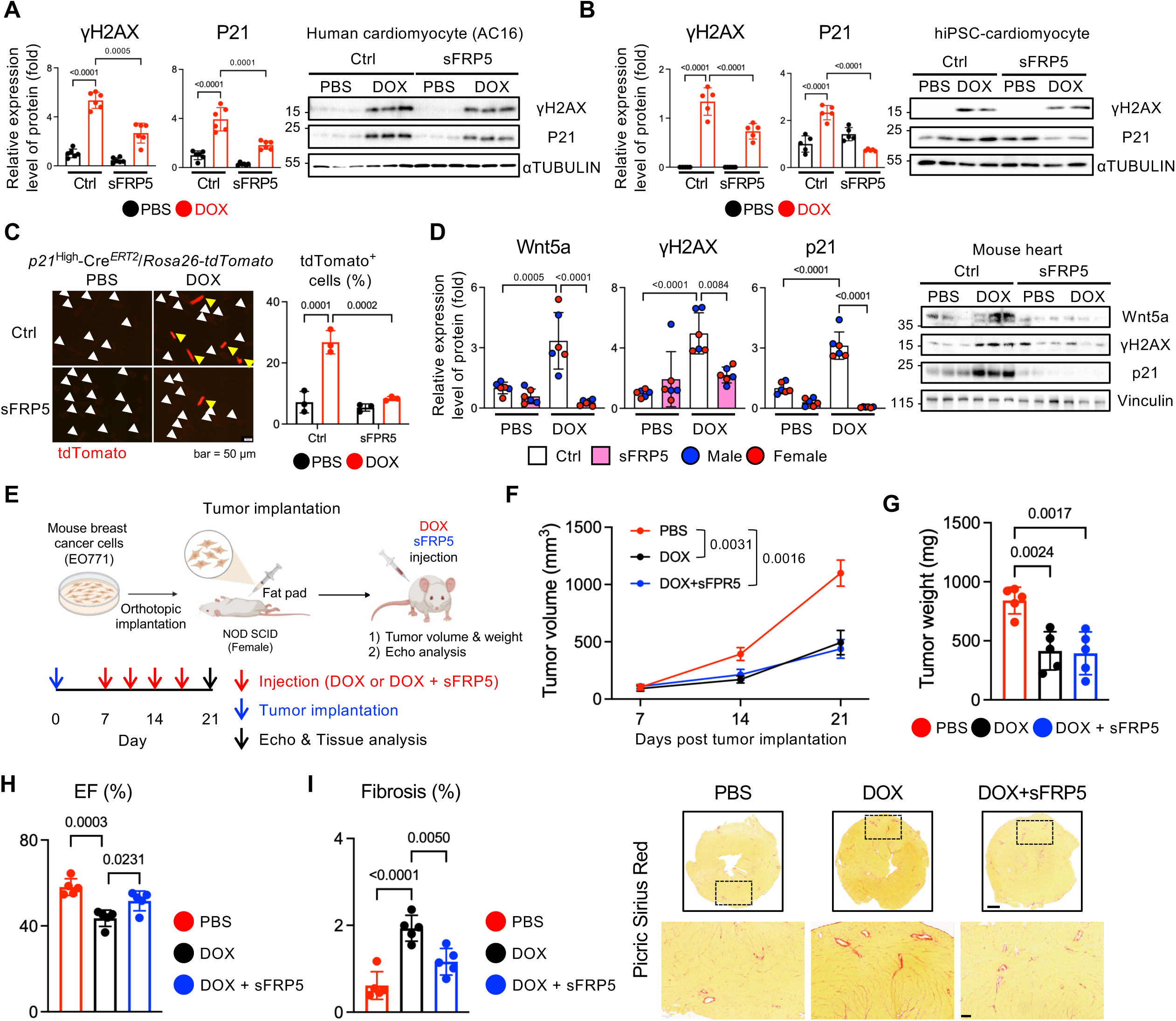
sFRP5 suppresses DOX-induced cardiomyocyte senescence and protects against cardiotoxicity without compromising anti-tumor efficacy. (**A** and **B**) AC16 cardiomyocytes and hiPSC-derived cardiomyocytes were treated with PBS or DOX (500 nM) with or without sFRP5 (400 ng/ml) for 48 hours. Cell lysates were subjected to Western blotting with anti-γH2AX, p21 and αTubulin antibodies. Protein levels were normalized by αTubulin. n = 6. (C) Representative fluorescence images and quantification of adult mouse cardiomyocytes from *p21*^High^-Cre^ERT2^;*Rosa26-tdTomato* mice and treated with PBS or DOX (100 nM) with or without sFRP5 (400 ng/ml) for 48 hours. White and yellow arrowheads indicate tdTomato-negative and tdTomato-positive *p21*^High^ cardiomyocytes, respectively. Scale bar: 50 μm. n = 3. (**D**) Wild-type mice were treated weekly with PBS or DOX (5 mg/kg, i.p.) with or without sFRP5 (20 μg/kg, i.p.) for 4 weeks. One week after the final injection, heart lysates were subjected to Western blotting with anti-Wnt5a, γH2AX, p21 and Vinculin antibodies. Protein levels were normalized by Vinculin. n = 6. (**E**-**I**) EO771 cells were orthotopically implanted into the mammary fat pads of female NOD SCID mice, followed by treatment with PBS, DOX (5 mg/kg, i.p.) or DOX plus sFRP5 (20 μg/kg, i.p.). (**E**) Experimental schematic. (**F**) Tumor volume (**F**) and final tumor weight (**G**) in EO771 tumor-bearing mice treated with PBS, DOX or DOX plus sFRP5. (**H**) LVEF assessed by echocardiography. n = 5. (**I**) Quantification and representative Picric Sirius Red staining of myocardial fibrosis. Scale bars: 500 μm (upper) and 100 μm (bottom). n = 5. Data are presented as mean ± SD. Statistical significance was determined by two-way ANOVA followed by Sidak’s multiple-comparison test (**A**, **B**, **D**) or one-way ANOVA followed by Tukey’s multiple-comparison test (**F**-**I**). For **F**, Final tumor volumes were compared. Exact P values are shown in the graphs.

We next asked whether sFRP5 alters the anticancer efficacy of DOX. *In vitro*, DOX reduced the cell viability of both mouse EO771 and human MDA-MB-231 breast cancer cells. Recombinant Wnt5a or sFRP5 alone did not significantly affect cancer cell viability and co-treatment with recombinant Wnt5a or sFRP5 did not alter the cytotoxic effect of DOX in either cell line (**Supplementary Figure 6B**). These results suggest that modulation of Wnt5a signaling does not interfere with DOX-induced cancer cell killing *in vitro*.

We next evaluated the effect of sFRP5 on DOX anti-tumor efficacy *in vivo* using an EO771 orthotopic breast cancer model. EO771 cells were implanted into the mammary fat pads of female mice, which were then treated with PBS, DOX or DOX plus sFRP5 (**Figure 6E**). DOX significantly suppressed tumor growth and reduced final tumor weight, and these anti-tumor effects were preserved even in the presence of sFRP5 (**Figure 6, F and G**). Importantly, in the same tumor-bearing mice, sFRP5 attenuated DOX-induced reductions in LV ejection fraction and increases in myocardial fibrosis, indicating that sFRP5 confers cardioprotection without a loss of DOX anti-tumor efficacy (**Figure 6, H and I**). Consistent with these findings, similar results were observed in a human MDA-MB-231 xenograft model, in which co-administration of sFRP5 did not diminish DOX-mediated tumor suppression (**Supplementary Figure S6, C and D**). Taken together, these findings indicate that sFRP5 protects against DOX-induced cardiac injury while preserving the anti-tumor activity of DOX in mouse models of breast cancer.

## Discussion

In this study, we identify Wnt5a as a central mediator of DOX-induced cardiotoxicity and uncover its mechanistic role in driving cardiomyocyte senescence and promoting maladaptive remodeling of the heart. The clinical relevance of this pathway is highlighted by our observation of elevated Wnt5a serum levels in DOX-treated cancer patients, as well as in human cardiomyocytes exposed to DOX. Using genetic and pharmacological approaches, we demonstrate that Wnt5a orchestrates a feed-forward signaling loop through its receptor Fzd2, engages the Pml-p53 axis to amplify the production of Wnt5a and propagates the senescent burden into neighboring fibroblasts. Furthermore, pharmacological inhibition of Wnt5a with sFRP5 protects against DOX-induced cardiomyopathy without compromising the chemotherapeutic efficacy of DOX in preclinical models of breast cancer. Together, these findings establish Wnt5a as a master regulator of DOX cardiotoxicity and highlight Wnt5a-Fzd2 pathway signaling as a promising therapeutic target.

### Cardiomyocyte-derived Wnt5a as the initiating trigger of DOX-induced cardiomyopathy

Our findings demonstrate that endogenous Wnt5a in cardiomyocytes is the critical initial determinant of DOX-induced senescence and cardiomyopathy. Following DOX treatment, Wnt5a protein levels rise sequentially, emerging first in cardiomyocytes before extending to non-myocyte populations. Crucially, genetic ablation of *Wnt5a* in cardiomyocytes also completely abolished *Wnt5a* mRNA expression in other non-myocyte cell populations in the early stage of DOX exposure, indicating that cardiomyocytes are the initiating source of pathogenic Wnt5a. By breaking this initial trigger in cardiomyocytes, we blocked senescence propagation and preserved cardiac function. The precise molecular mechanisms through which cardiomyocytes, rather than other cell types, rapidly and preferentially upregulate Wnt5a transcription in response to DOX remain an area for future investigation. Given that the Pml-p53 axis promotes the expression of SASP genes alongside Wnt5a, we hypothesize that Wnt5a transcription is rapidly amplified through a positive feedback mechanism involving the Pml-p53 pathway within the stressed myocardium.

### Mechanism by which Wnt5a induces cardiomyocyte senescence

A key question arising from our study is how Wnt5a drives senescence in cardiomyocytes and other cardiac cell types. As a non-canonical Wnt ligand, Wnt5a can trigger senescence via multiple mechanisms in a context-dependent manner. In our model, DOX-induced Wnt5a upregulation temporally coincided with increased senescence phenotypes, including elevated p21 and γH2AX. Previous studies suggest that Wnt5a can facilitate cell cycle arrest and senescence by antagonizing canonical Wnt signaling in ovarian cells and tendon stem/progenitor cells, or by activating the JAK-STAT pathway to increase p16 expression and SASP secretion in tendon stem/progenitor cells (38, 39). Under genotoxic stress, Wnt5a secretion can also create an ATR/ATM-dependent positive feedback loop that reinforces the DNA damage response and increases p21 (40). This mechanism likely explains how Wnt5a not only reinforces senescence in cardiomyocytes (autocrine effect) but also spreads it to neighboring fibroblasts and other non-myocytes (paracrine effect), amplifying the senescent burden across the heart.

Our inhibitor analysis implicates p38 MAPK as a potential downstream mediator of Wnt5a-induced senescence marker induction in cardiomyocytes. Together with previous evidence that p38 MAPK inhibition attenuates DOX-induced cardiotoxicity *in vivo* (41), our findings raise the possibility that Wnt5a-Fzd2 signaling may serve as an upstream trigger of p38 MAPK-dependent cardiotoxic signaling. Further studies will be needed to directly test whether Wnt5a-Fzd2 activation drives p38 MAPK signaling *in vivo* during DOX-induced injury.

Another important question is whether DOX-induced senescence produces a secretory phenotype that is distinct from aging-associated cardiomyocyte senescence. Atypical SASP factors, including Gdf15, Edn3 and Tgfb2, have been linked to aged cardiomyocytes (17). However, in our DOX model, only *Gdf15* was induced and was reduced by cardiomyocyte *Wnt5a* deletion. This pattern suggests that DOX-induced cardiomyocyte senescence may not simply recapitulate aging-associated senescence but instead may generate a context-specific SASP program shaped by genotoxic stress and Wnt5a signaling. Defining how Wnt5a controls distinct SASPs across DOX injury, aging and other cardiac stress conditions will be an important area for future investigation.

### The Wnt5a-Fzd2 feed-forward loop and paracrine spread of senescence

A novel aspect of our study is the identification of a feed-forward Wnt5a-Fzd2 loop that propagates senescence across cardiac cell types. While cardiomyocytes act as the primary initiators of Wnt5a expression after DOX treatment, steady chronic increases in Wnt5a expression in the heart appear to also be mediated by non-myocyte populations. In particular, we found that cardiac fibroblasts are highly responsive to paracrine Wnt5a. Our scRNA-seq analyses showed that although Wnt5a-Fzd2 signaling is prominent as a cardiomyocyte autocrine loop at day 1 after DOX treatment, cardiomyocyte-to-fibroblast/stromal cell communication becomes strong and fibroblast/stromal cell autocrine signaling is established by week 2. Furthermore, specific deletion of the Wnt5a receptor, Fzd2, in cardiac fibroblasts reduced global cardiac senescence and preserved function to a degree comparable to the cardiomyocyte-specific Wnt5a KO model. These results position Wnt5a not merely as a stress-induced signal but as a self-amplifying driver of maladaptive remodeling. Thus, besides preventing the initial induction of Wnt5a in cardiomyocytes, the amplification mechanisms, including Fzd2 in fibroblasts and production of Wnt5a in senescent cells in non-myocyte populations, may be targeted for cardioprotection. Further clarification of the involvement of senescent cells of various cell types in the amplification of Wnt5a in the DOX-treated heart and their spatio-temporal regulation should lead to the development of improved interventions for DOX-induced cardiomyopathy.

### Translational relevance of targeting the Wnt5a-Fzd2 axis

From a translational perspective, our findings suggest that Wnt5a-Fzd2 signaling may have both biomarker and therapeutic relevance in anthracycline-associated cardiotoxicity. The increase in circulating WNT5A after anthracycline therapy suggests that WNT5A may have potential as a blood-based marker of early cardiac stress during cancer treatment. Further studies in larger perspective cohorts will be needed to determine whether circulating WNT5A predicts subsequent cardiac dysfunction or treatment-related cardiotoxicity.

Therapeutically, our findings identify use of the soluble decoy receptor sFRP5 as a highly translational strategy for mitigating chemotherapy-induced cardiomyopathy. Systemic administration of sFRP5 abrogated DOX-induced cardiac dysfunction and fibrosis while reducing cardiomyocyte senescence. Notably, sFRP5 did not interfere with the anti-cancer activity of DOX in breast cancer xenograft models. This selectivity is a critical advance, as most cardioprotective interventions risk blunting the efficacy of chemotherapeutic agents (42, 43). By specifically uncoupling Wnt5a-driven cardiac senescence from DOX-mediated tumor apoptosis, sFRP5 provides a safe therapeutic window. Beyond directly antagonizing Wnt5a, sFRP5 may exert additional benefits by reducing pro-inflammatory cytokine production and improving metabolic regulation, as previously reported in other metabolic and fibrotic diseases (44–48).

### Limitations and future directions

Several limitations warrant discussion. First, although sFRP5 is recognized as a competitive antagonist for Wnt5a, it can also interact with and antagonize other Wnt family ligands, including Wnt3a and Wnt11 (49, 50). While we focused on the Wnt5a-Fzd2 axis, it remains possible that other non-canonical Wnt ligands redundantly contribute to DOX cardiotoxicity. Second, our *in vivo* cancer experiments utilized a breast tumor model. Since DOX is a mainstay therapy for leukemias and sarcomas, validating sFPR5 efficacy across diverse tumor models will be essential for broad clinical generalization. Third, to more closely model therapeutic applicability in patients, it will be critical to investigate whether delayed administration of sFRP5 after the onset of DOX-induced cardiac injury still confers substantial cardioprotection. Finally, long-term survival studies are needed to determine whether sFRP5 provides durable protection, or whether combinatorial strategies utilizing senolytics and dexrazoxane, an FDA-approved medication for DOX-induced cardiomyopathy, could further enhance cardiovascular outcomes in cancer survivors.

## Methods

### Sex as a biological variable

Sex was not considered as a biological variable.

### Human serum samples

Paired serum samples collected before initiation of anthracycline therapy and at an approximately 3-month follow-up after treatment were obtained from patients with breast cancer. Samples were provided by Dr. Sebastiano Sciarretta (Sapienza University of Rome) under an approved human subjects protocol (PROT.0157/2025). All samples were de-identified before analysis.

### Enzyme-linked immunosorbent assay (ELISA)

The human WNT5A ELISA kit was purchased from CUSA BIO (CSB-EL026138HU). Concentrations of WNT5A in human serum samples were measured by ELISA according to the manufacturer’s protocol.

### Cell culture

The human cardiomyocyte cell line AC16 was purchased from PromoCell (C-128010) and maintained according to the manufacturer’s instructions. hiPSC-derived cardiomyocyte was purchased from FUJIFILM (01434) and maintained according to the manufacturer’s instructions.

The mouse breast cancer cell line EO771 was kindly provided by Dr. Raymond Birge (Rutgers NJMS) and cultured in DMEM supplemented with 10% FBS. The human breast cancer cell line MDA-MB-231 was kindly provided by Dr. Lai-Hua Xie (Rutgers NJMS) and maintained in RPMI 1640 (11879020, Thermo Fisher Scientific) supplemented with 10% FBS. The human breast cancer cell MCF7 was purchased from Millipore Sigma (86012803-1VL) and cultured in MEM (M5650, Millipore Sigma) supplemented with human insulin (4 mg/ml; 1258014, Thermo Fisher Scientific), L-Glutamine (1:100; 25030081, Thermo Fisher Scientific) and 10% FBS.

### Mice

*Wnt5a*^fl/fl^ mice (#26626), *Myh6*-Cre mice (#009074) and *Rosa-CAG-LSL-tdTomato* mice (#007914) were purchased from the Jackson Laboratory and maintained in our mouse facility. *Pml*^fl/fl^ mice (#T051860) were purchased from GemPharmatech Co., Ltd. Cardiac-specific *Wnt5a* homozygous knockout mice (*Wnt5a*^cKO^) were generated by crossing *Wnt5a*^fl/fl^ mice with *Myh6*-Cre mice. The *p21*^High^-Cre^ERT2^ mice and *Fzd2*^fl/+^ mice were generated by the Genome Editing Shared Resource (GESR) at Rutgers, The State University of New Jersey, using previously published designs (32, 51). *p21*^High^-Cre^ERT2^ mice were crossed with *Wnt5a*^fl/fl^ mice or *Rosa-CAG-LSL-tdTomato* (hereafter *Rosa-tdTomato*) mice to generate *p21*^High^-Cre^ERT2^/*Wnt5a*^fl/+^ and *p21*^High^-Cre^ERT2^/*Rosa-tdTomato* mice, respectively. *Tcf21*-Cre^ERT2^ mice were kindly provided by Dr. Dominic Del Re (Rutgers New Jersey Medical School). Cardiac-specific *Fzd2* heterozygous mice (*Fzd2*^chKO^) and cardiac fibroblast-specific *Fzd2* heterozygous mice (*Fzd2*^cfhKO^) were obtained by crossing *Fzd2*^fl/+^ mice with *Myh6*-Cre mice or *Tcf21*-Cre^ERT2^ mice, respectively. Cardiac-specific *Pml* heterozygous knockout mice (*Pml*^chKO^) were generated by crossing *Pml*^fl/fl^ mice with *Myh6*-Cre mice. All animal experiments were approved by the Institutional Animal Care and Use Committee of Rutgers New Jersey Medical School. Both male and female mice were used.

### Doxorubicin administration to mice

Doxorubicin (DOX; D1515-10MG, Sigma-Aldrich) was administered to mice via intraperitoneal injection at a dose of 5 mg/kg (dissolved in PBS) once a week for 4 consecutive weeks (52). Mice injected with PBS were used as controls.

### Isolation of adult mouse cardiomyocytes

For primary culture, adult mouse cardiomyocytes (AMCMs) were isolated as previously described with minor modification (53). Briefly, hearts were perfused with 7 ml EDTA buffer via the right ventricle using a 27-gauge needle to arrest cardiac contractions. After clamping the ascending aorta, enzymatic digestion was performed by sequential injection of 10 ml EDTA buffer and 3 ml perfusion buffer followed by 20 ml perfusion buffer containing collagenase type II (2.5 mg/ml, Worthington) through the LV. Digestion was terminated by adding 5 ml perfusion buffer supplemented with 5% FBS. Cardiomyocytes and non-cardiomyocytes were separated by four sequential rounds of gravity sedimentation with gradual addition of perfusion buffer and culture medium (M199, M4530, Millipore Sigma) supplemented with 0.1% BSA, 10 mM 2,3-butanedione monoxime (BDM; Millipore Sigma) and 1% chemically defined lipid supplement (11905031, Thermo Fisher Scientific). Cardiomyocytes were plated onto laminin (5 µg/ml)-coated cell culture dishes for subsequent experiments.

For scRNA-seq, AMCMs were enzymatically isolated using a Langendorff perfusion system as previously described (54). Briefly, mice were deeply anesthetized with isoflurane. Hearts were rapidly excised, cannulated via the aorta and perfused retrogradely at 37°C with a Ca²⁺-free Tyrode’s solution (136 mM NaCl, 5.4 mM KCl, 1 mM MgCl₂, 0.33 mM NaH₂PO₄, 10 mM glucose, and 10 mM HEPES; pH 7.4) containing 0.2 mg/mL Liberase TH Research Grade (Sigma-Aldrich) for 10 to 12 minutes. Following enzymatic digestion, the hearts were removed from the Langendorff apparatus and transferred to a petri dish, where the left ventricles were gently mechanically dissociated using fine forceps. The resulting cell suspension was filtered through a 200-μm strainer to remove undigested tissue. Freshly isolated cardiomyocytes were immediately subjected to fixation for subsequent scRNA-seq.

### scRNA-seq analyses

DOX (5 mg/kg) was administered intraperitoneally according to **Figure 2D**. Control mice were injected with PBS. Adult mouse heart cells were isolated from both PBS-treated (n = 4), DOX-treated Day 1 (n = 4) and DOX-treated Week 2 (n = 4) mice. Freshly isolated cardiomyocytes and non-myocytes were immediately fixed using Evercode Fixation v2 kits (Parse Biosciences), following the manufacturer’s instructions. Cells were processed to sublibraries using the Evercode WT v2 kit (Parse Biosciences). Barcoding and library generation were performed according to the manufacture’s protocols.

Sequencing reads were aligned to the GRCm38 mouse reference genome using the Parse Biosciences data processing pipeline. Downstream analyses were performed using Scanpy in Python. Following strict quality control to exclude low-quality cells and doublets, raw count matrices were normalized (10,000 per cell), long-transformed and clustered via the Leiden algorithm following PCA and UMAP dimensionality reduction. Major cell types were annotated using established canonical marker genes.

To investigate the senescence-associated stress response specifically within the cardiomyocyte fraction, cells were dichotomized into positive (*Cdkn1a*^+^) or negative (*Cdkn1a*^-^) subpopulations based on the detection of raw gene expression (> 0). The fraction of *Wnt5a*^+^ cells was calculated within these respective subpopulations. Furthermore, the expression dynamics of specific p53 downstream target genes across the *Cdkn1a*^+^ and *Cdkn1a*^-^ cardiomyocytes were evaluated and visualized using scaled dot plots.

To predict the strength of targeted intercellular paracrine signaling, a mass-action-based interaction score for the Wnt5a-Fzd2 ligand-receptor pair was computed. The interaction potential was calculated by multiplying the mean expression level of the ligand (*Wnt5a*) in the sender cell populations by the mean expression level of the receptor (*Fzd2*) in the recipient cell population. The calculated relative interaction scores were utilized to evaluate the paracrine signaling burden across experimental groups.

### Picrosirius Red staining

Hearts were fixed in 4% paraformaldehyde (PFA), embedded in paraffin and sectioned at 10 µm thickness. Myocardial fibrosis was assessed by Picrosirius Red staining as described previously (55). Fibrotic area was quantified as the percentage of collagen-stained (red) area relative to total myocardial area using ImageJ software.

### Immunofluorescence staining

AMCMs were cultured on 3.5 cm dishes containing coverslips coated with laminin (5 μg/ml). Cells were fixed with 4% PFA, permeabilized with 0.3% Triton X-100 in PBS, and blocked with 3% BSA in PBS for 90 minutes at room temperature. Cells were incubated overnight at 4 °C with primary antibodies, followed by 1 hour incubation with Alexa Fluor 488-, Alexa Fluor 594– or Alexa Fluor 648-conjugated secondary antibodies (Invitrogen). Coverslips were washed and mounted on glass slides using VECTASHIELD mounting medium with DAPI (Vector Laboratories). Fluorescence images were acquired using a confocal (Nikon) or epifluorescence microscope (Nikon).

### Adeno-associated virus

AAV9 vectors encoding either *cTNT*-*WNT5A* (AAV9-*cTNT*-*WNT5A*) or control GFP (AAV9-*cTNT*-*GFP*) were purchased from Vector Builder. C57BL/6J mice were administered a single dose of 1 × 10^11^ vg/mouse via intrajugular vein.

### Bulk RNA-sequencing

Total RNA was isolated from mouse hearts one week after AAV9-*cTNT*-*WNT5A* or AAV9-*cTNT*-*GFP* injection and sent to Novogene Corporation Inc. for RNA sequencing. RNA integrity was evaluated using an Agilent Technologies 2100 Bioanalyzer. Sequencing libraries were constructed using the NEBNext Ultra RNA Library Prep Kit for Illumina via oligo-(dT) magnetic bead mRNA purification, fragmentation and cDNA synthesis. Paired-end sequencing was performed on an Illumina NovaSeq 6000 platform. For bioinformatics analysis, adapter and low-quality reads were removed using fastp. Clean reads were mapped to the GRCm38 reference genome using HISAT2 and quantified with featureCounts. Differentially expressed genes (DEGs) were identified using the DESeq2 R package with log2 (fold change) > 1 or log2 (fold change) < –1 and adjusted p-value < 0.05.

### Chromatin immunoprecipitation sequencing (ChIP-seq)

ChIP-seq was performed according to the manufacturer’s protocol (ActiveMotif^®^) as described previously (56). Briefly, heart tissues from C57BL/6J wild-type mice treated with PBS (n = 4) or DOX (10 mg/kg, a single dose; n = 4) were harvested one week after the injection and crosslinked with 1% formaldehyde for 15 minutes, followed by quenching with glycerin for 5 minutes at room temperature. Samples were pooled, washed twice with cold PBS, lysed, and subjected to sonication to shear chromatin. Protein-DNA complexes were immunoprecipitated using an anti-p53 antibody (BS-8687R, Bios USA), followed by sequential washes and protein digestion with proteinase K. Reverse crosslinking was carried out at 65 °C and DNA was purified for downstream analysis. ChIPSeq libraries were prepared using the TruSeq ChIP Library Preparation Kit (Illumina) and sequenced on an Illumina Genome Analyzer 2. Sequence reads were aligned to the reference genome using Bowtie, and peak calling and annotation were performed using the Bioconductor ChIP-seq package. Motif enrichment analysis was conducted with HOMER, and peak visualization was carried out using the USC IGV browser.

### Tumor model for mouse

The orthotopic breast tumor model was established as previously described (57, 58). Mouse breast cancer cells (EO771, 1 × 10^5^ cells) or human breast cancer cells (MDA-MB-231, 1 × 10^6^ cells) were suspended in 50 μl of serum-free media with 50 μl of Matrigel (354230, Corning). The cell suspension was injected into the right inferior mammary fat pad of 8-week-old female NOD SCID mice under anesthesia. Tumor volume was measured using calipers and calculated as 0.5 X (long axis) X (short axis)^2^. Once tumors reached a volume of 50-100 mm^3^, mice were randomly assigned to treatment groups to ensure comparable mean tumor volumes at the time of randomization. Treatments were administered interperitoneally. After echocardiographic analysis, the mice were euthanized and tumors were excised and weighed one week after the final injection.

### Statistical analysis

All data points were obtained from independent biological samples unless otherwise indicated. In bar graphs, individual data values (*n*) are represented by symbols. Continuous data are expressed as mean ± SD. Comparisons between two groups were performed using two-tailed unpaired Student’s *t-*test or Wilcoxon matched-pairs signed-rank test, as appropriate. Comparisons among 3 or more groups were performed using one-way or two-way ANOVA followed by Tukey’s or Sidak’s multiple comparison test, as indicated in the figure legends. Categorical variables were compared using Fisher’s exact test. A P value of < 0.05 was considered statistically significant. Detailed statistical parameters are provided in the figure legends. Statistical analyses and graph generation were performed using GraphPad Prism 10 (GraphPad software).

### Study approval

All animal procedures were reviewed and approved by the IACUC of Rutgers Biomedical and Health Sciences, Newark, New Jersey, USA.

### Data availability

scRNA-seq data and ChIP-seq data are deposited in the National Center for Biotechnology Information Gene Expression Omnibus (NCBI GEO). Published human snRNA-seq data are available in the GEO database (GSE292067). Supporting data values are provided in Supplemental Data.

### Author contributions

E-A. S. and J.S. designed the study. E-A. S. performed most of the experiments, analyzed data and interpreted results. PZ conducted the *in vitro* and *in vivo* experiments. S.I., M.M., K.T. and T.T. conducted echocardiographic measurements of mice. T.T. cultured hiPSC-derived cardiomyocytes. N.F. and L-H. X. isolated adult mouse cardiomyocytes for scRNA-seq. E-A. S. obtained and analyzed scRNA-seq data. G.Y. and P.R. generated genetically altered mouse lines. V.G and R.B. provided expertise in establishing the mouse tumor model. D.V., M.F., V.V. and S.S. provided human serum samples and clinical data. E-A. S. and J.S wrote the paper. J.S. oversaw the entire study and generated project resources. All authors reviewed and commented on the manuscript.

### Funding support

This study was supported in part by U.S. Public Health Service grants HL91469, HL112330, HL138720, HL144626, and HL150881 (J.S.). This work was also supported by an American Heart Association Predoctoral Fellowship 915784 (E-A. S.), Merit Award 20 Merit 35120374 (J.S.), Transformational Project Award 25TPA1481361 (J.S.), and by the Foundation Leducq Transatlantic Network of Excellence 15CVD04 (J.S.).

## Supporting information

Individual data

## Acknowledgements

We thank Daniela Zablocki for critical reading of the manuscript.

---

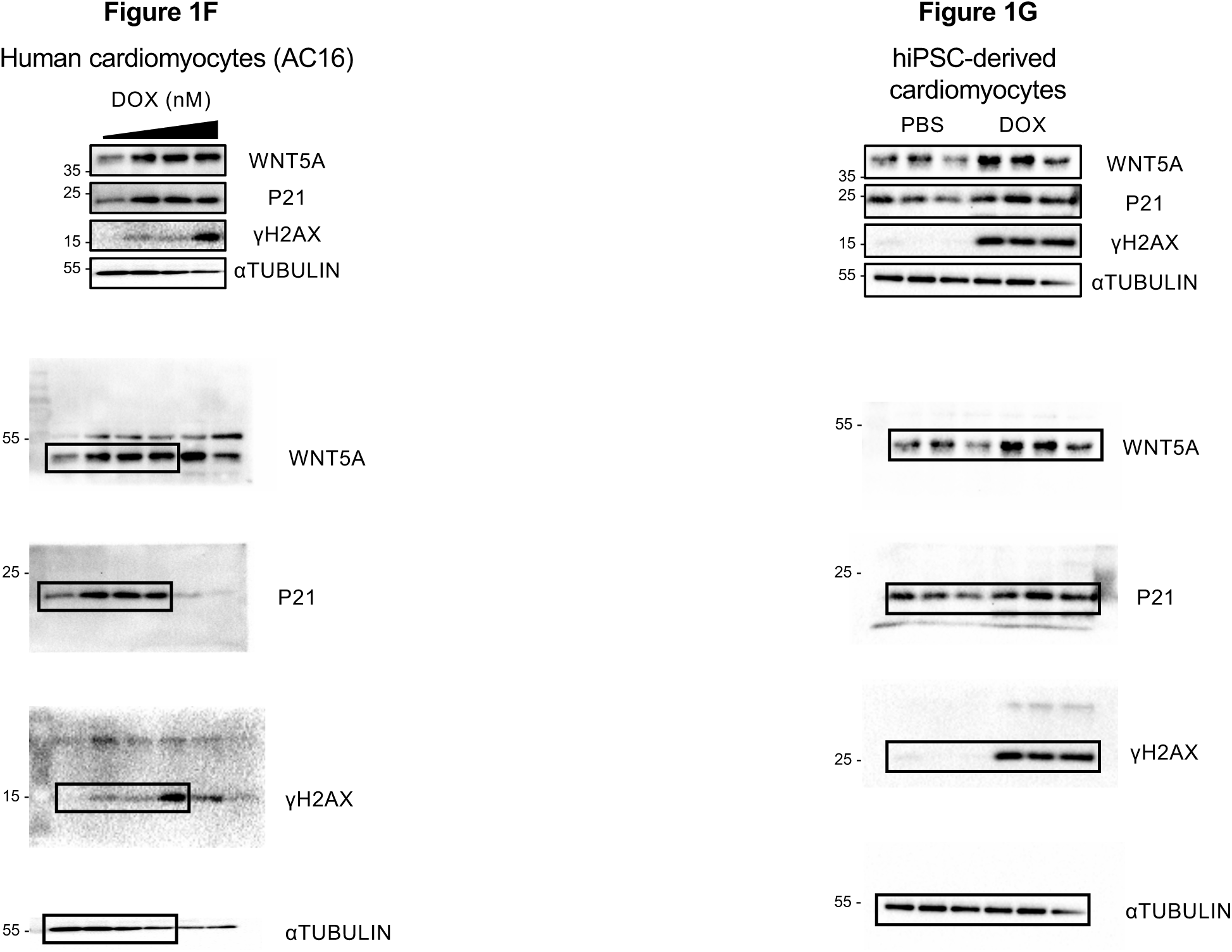

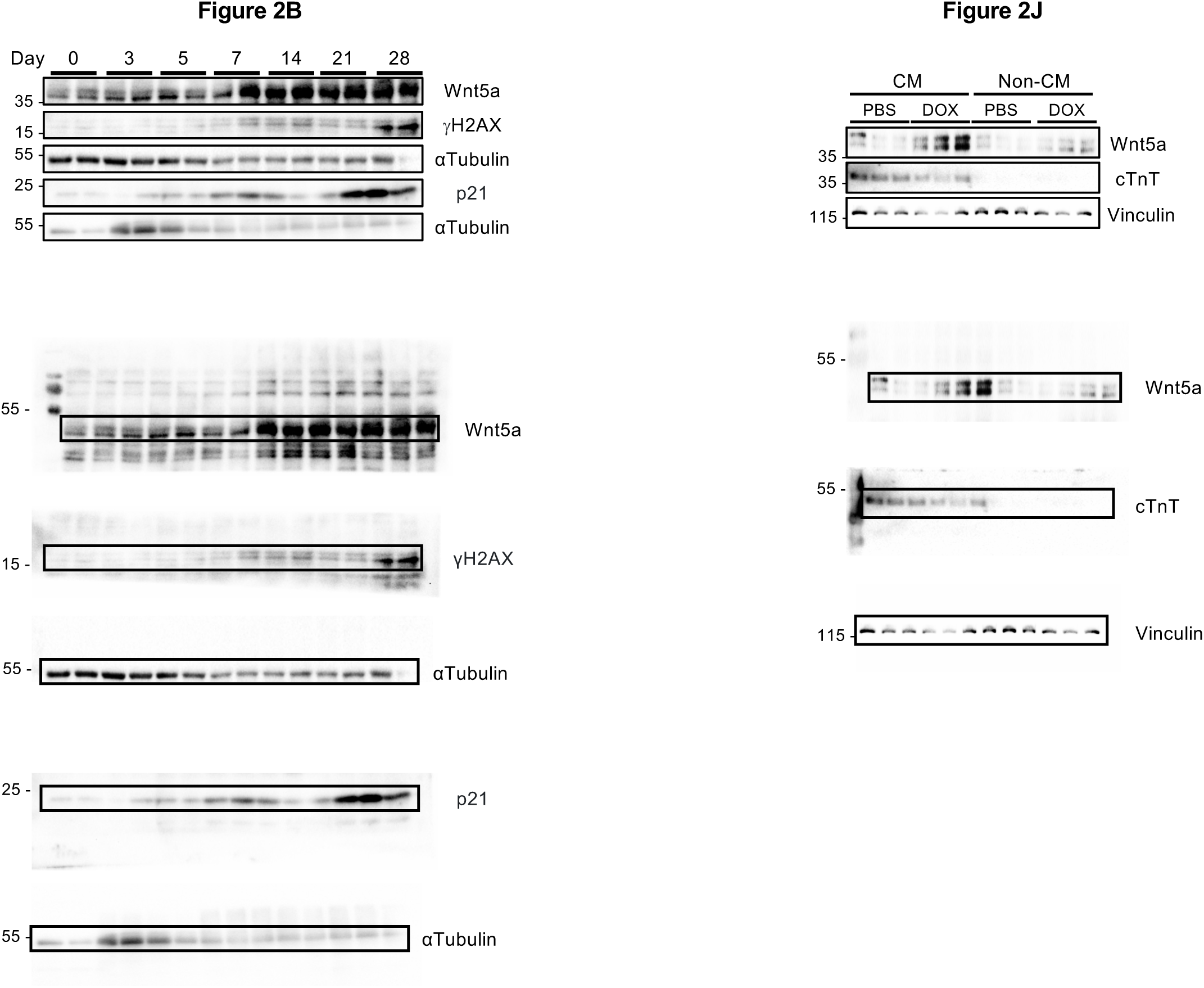

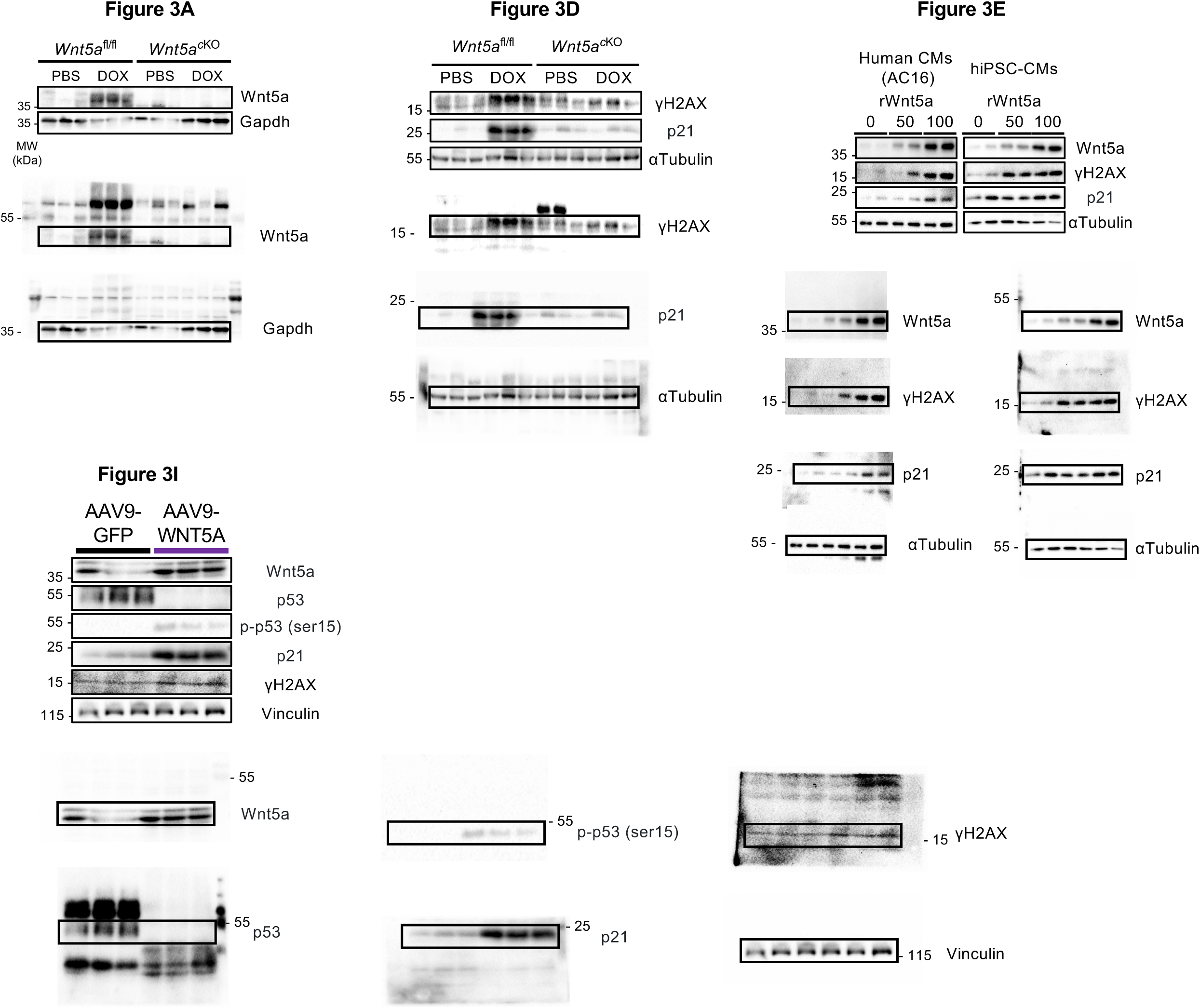

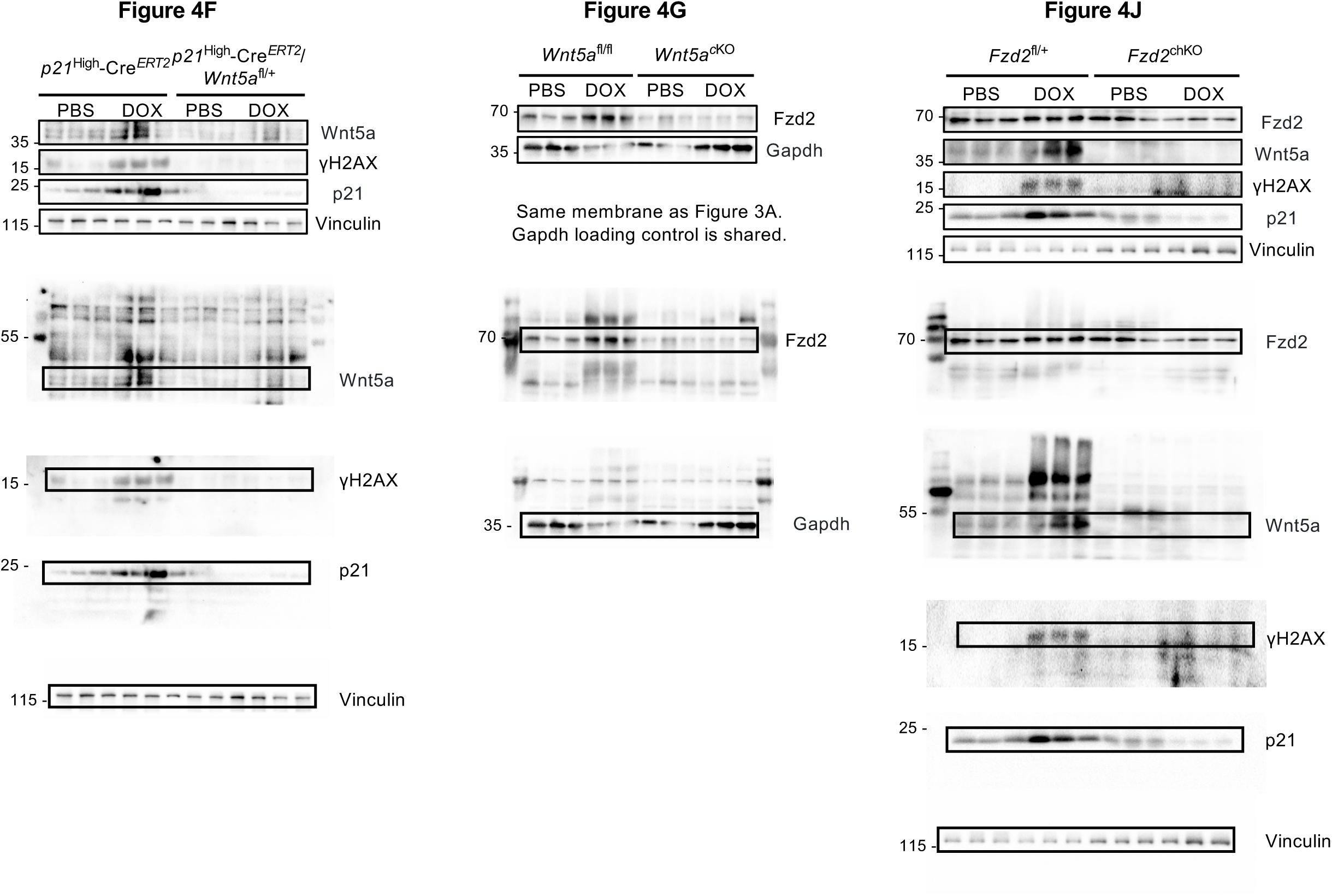

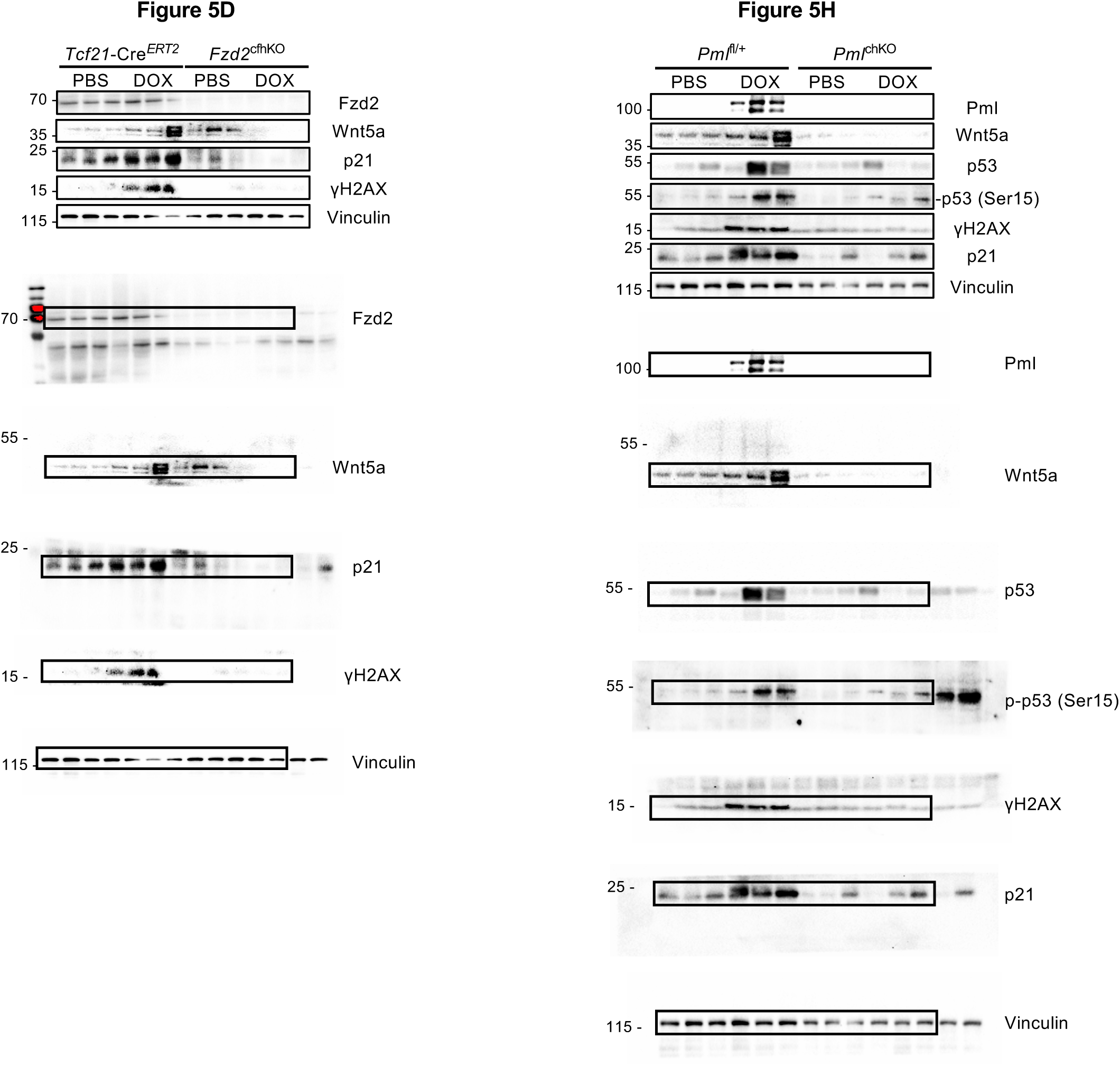

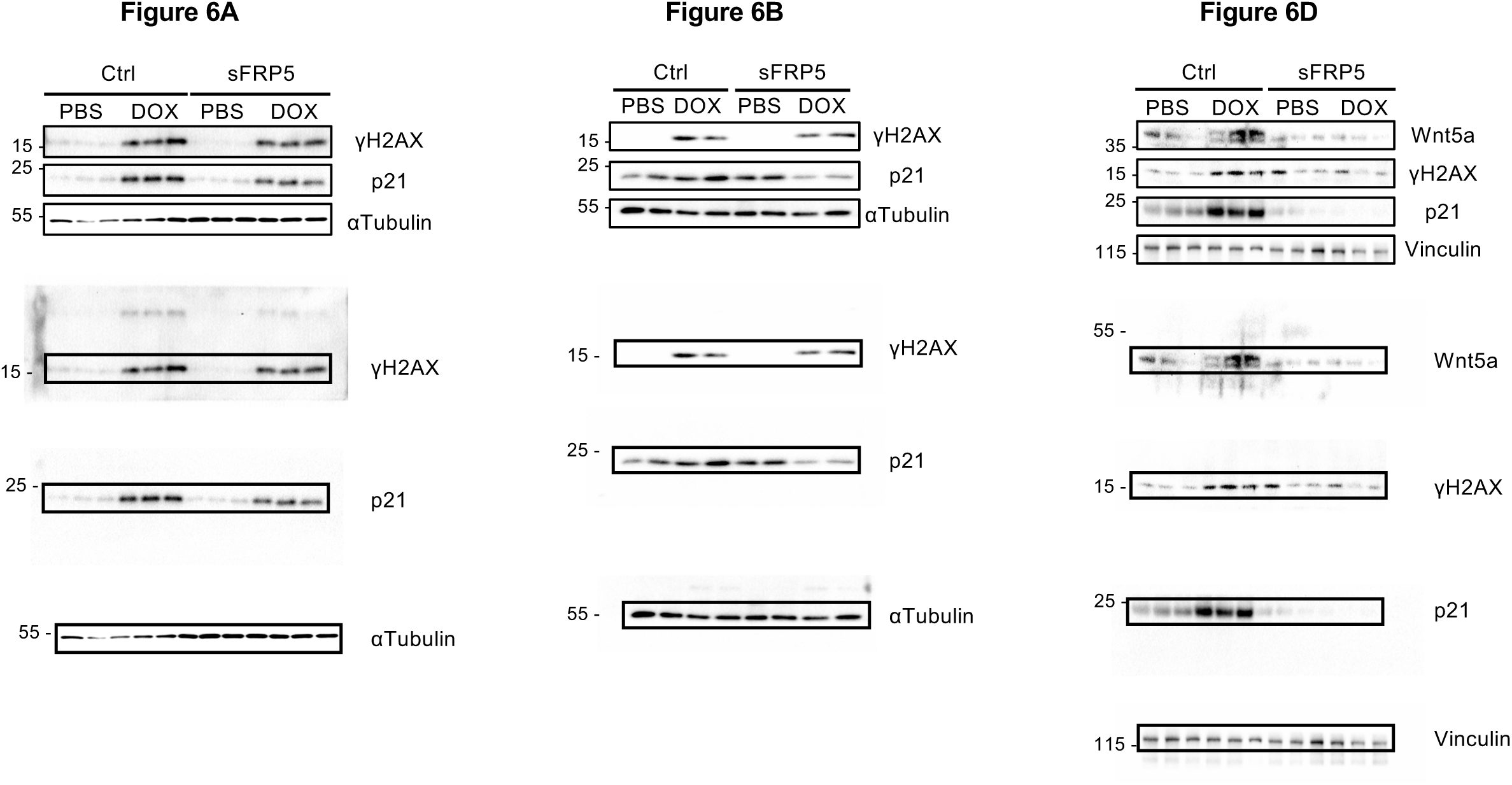

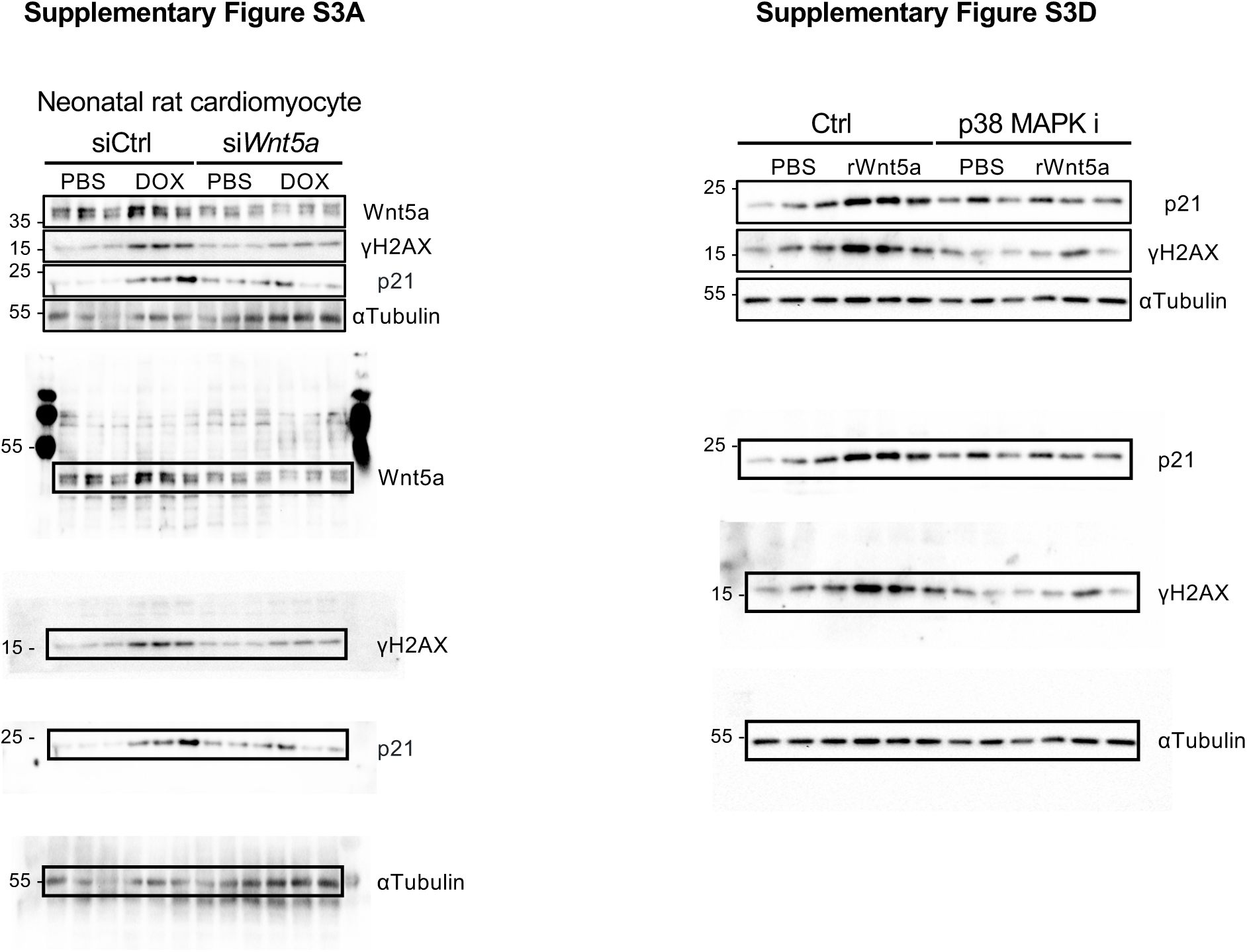

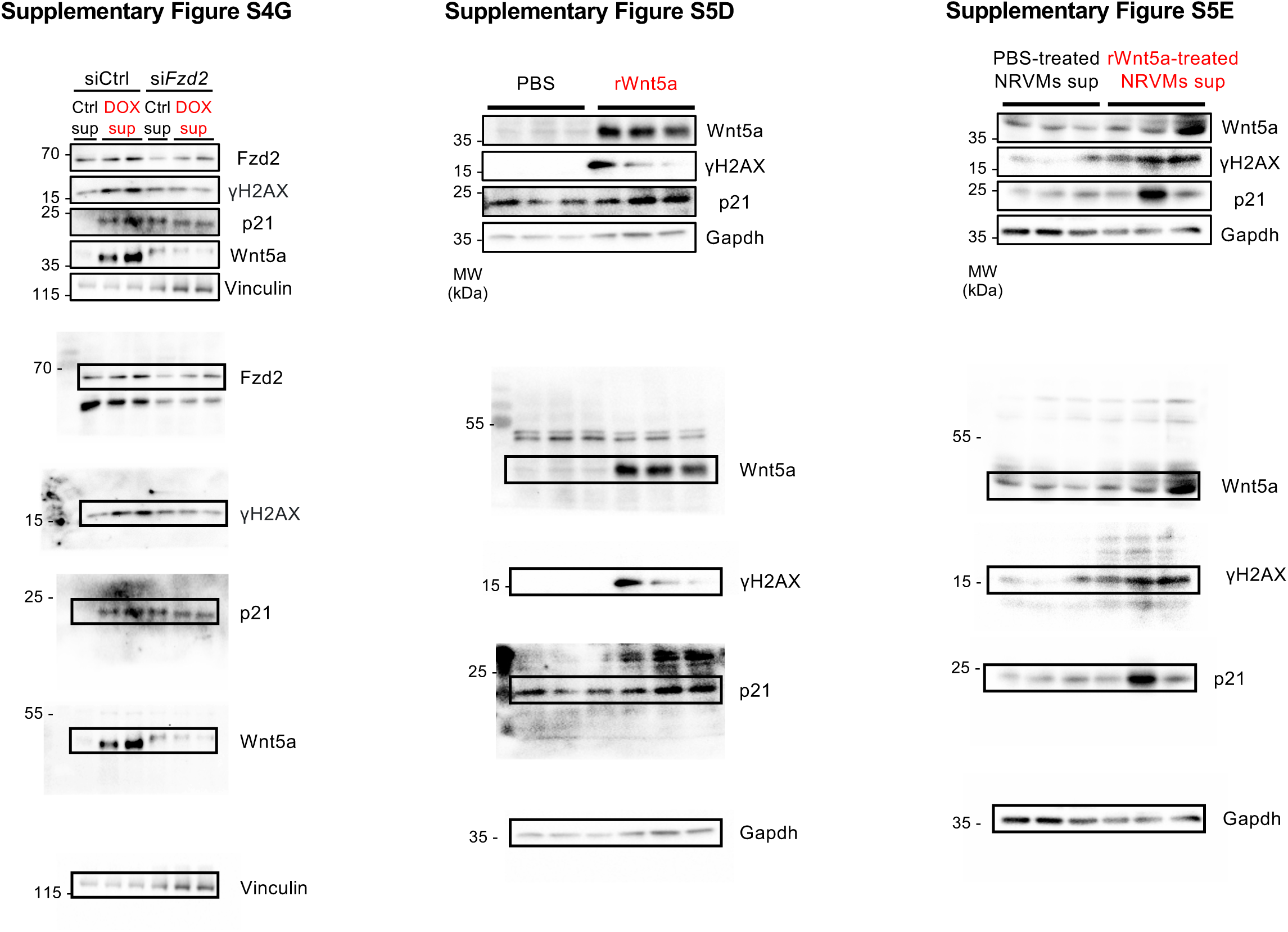

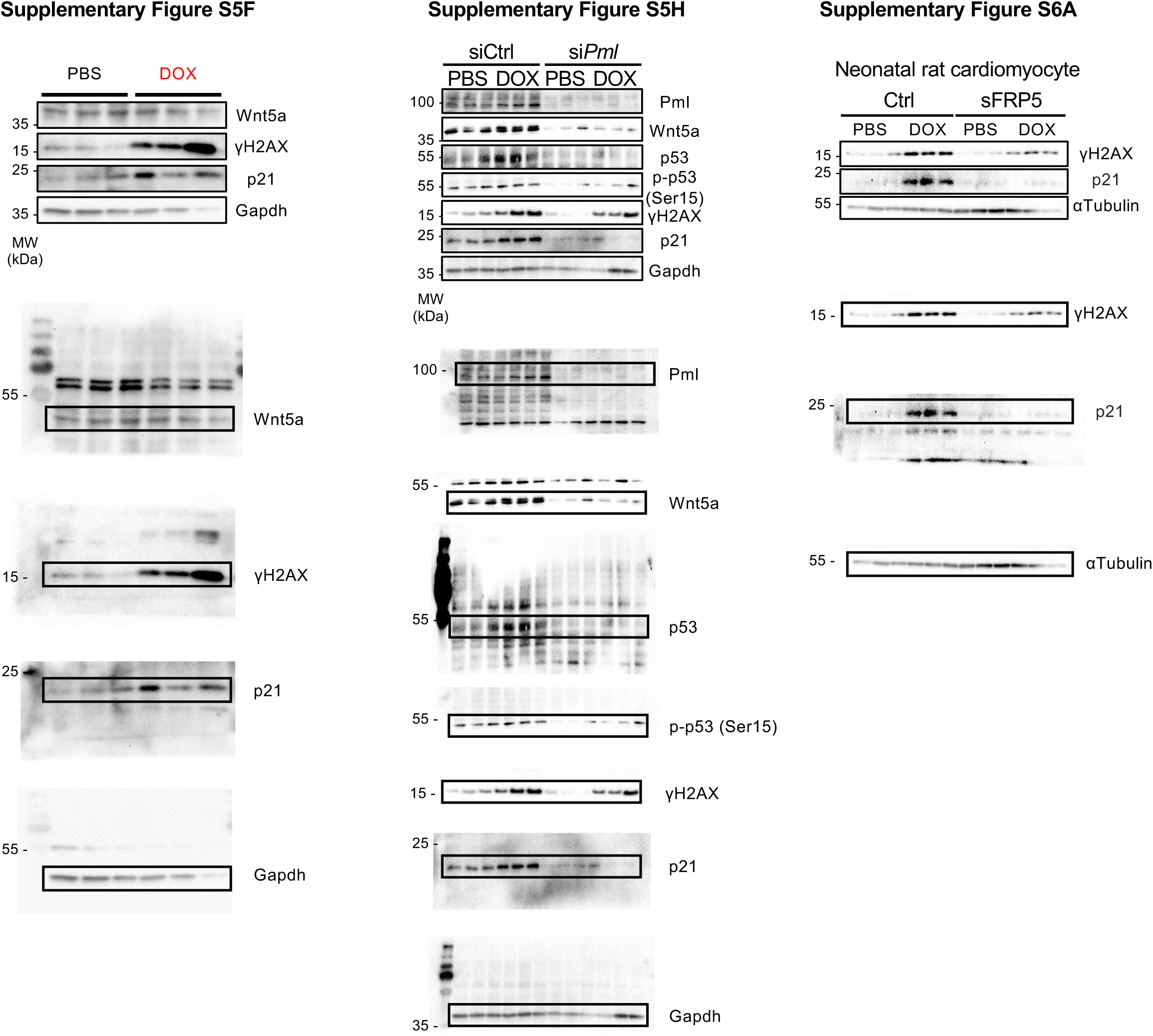

