## Supplemental methods and Figures for "Cardiomyocyte-derived Wnt5a drives doxorubicin-induced cardiomyopathy by amplifying cellular senescence"

Cardiovascular Research Institute, Rutgers Biomedical and Health Sciences, 185 South Orange Ave, MSB G609, Newark, NJ 07103

### Supplementary Methods

#### Primary culture of neonatal rat ventricular cardiomyocytes

Neonatal rat ventricular cardiomyocytes (NRVMs) and neonatal rat cardiac fibroblasts (NRCFs) were isolated as described previously (53). Briefly, neonatal rats (postnatal day 1, Wistar Institute (WI) BR-Wistar rats, Harlan Laboratories) were deeply anesthetized with 5% isoflurane inhalation (5-10 minutes), after which euthanasia was carried out by rapidly excising the hearts. A cardiomyocyte-enriched fraction and cardiac fibroblast-enriched fraction were obtained by centrifugation through a discontinuous Percoll gradient. NRVMs were cultured in complete medium containing DMEM/F-12 supplemented with 5% horse serum, 4 mg/ml transferrin, 0.7 ng/ml sodium selenite, 2 g/l bovine serum albumin (fraction V), 3 mM pyruvate, 15 mM HEPES (pH 7.1), 100 mM ascorbate, 100 mg/l ampicillin, 5 mg/l linoleic acid and 100 mM 5-bromo-2′-deoxyuridine. NRCFs were cultured in DMEM (11960044, Thermo Fisher Scientific) supplemented with 10% FBS (100-106, GeminiBio). Culture dishes were coated with 0.3% gelatin.

#### Echocardiography

Transthoracic echocardiography was performed under continuous isoflurane inhalation general anesthesia (3% induction, 1-2% maintenance) using a high-resolution Micro-Ultrasound system (Vevo 3100, FUJIFILM VisualSonics Inc., Toronto, Canada). Two-dimension (short-axis)-guided M-mode measurements were acquired at the level of the papillary muscles. Measurements of the LV internal diameter were obtained from three or more cardiac cycles and averaged. LV end-diastolic dimension (LVIDd) was measured at the point of maximal LV diastolic dimension, and LV end-systolic dimension (LVIDs) was measured at the point of the most anterior systolic excursion of the posterior wall. LV ejection fraction (LVEF) was analyzed using Vevo LAB software (FUJIFILM VisualSonics Inc.) to evaluate systolic function.

#### RT-qPCR

Total RNA was extracted from mouse heart tissues or cultured cells using TRIzol reagent (15596026, Thermo Fisher Scientific). After homogenizing the cell samples with TRIzol reagent, RNA purification was performed using the RNeasy Plus Mini Kit (74136, QIAGEN) according to the manufacturer's instructions. RNA concentration was measured using a NanoDrop spectrophotometer (Thermo Fisher Scientific). Reverse transcription of total RNA was performed using the RNA to cDNA EcoDry Premix kit (cat no. 639548, Takara Bio). qPCR was performed using PowerUp SYBR Green Master Mix (A25742, Thermo Fisher Scientific). The mRNA levels of target genes were normalized by comparison to the mRNA level of *Tbp* control using the $2^{-\Delta\Delta Ct}$ method. Primers used for qPCR are listed below.

*Wnt5a*, ACACAACAATGAAGCAGGCCGTAG and GGAGTTGAAGCGGCTGTTGACC;

*Fzd1*, CAAGGTTTACGGGCTCATGT and TGAACAGCCGGACAGGAAAA;

*Fzd2*, CCGACGGCTCTATGTTCTTC and TAGCAGCCGGACAGAAAGAT;

*Fzd3*, TGGGTTGGAAGCAAAAAGAC and CCTGCTTTGCTTCTTTGGTC;

*Fzd4*, GCCAATGTGCACAGAGAAGA and AGGTGGTGGAGATGAAGCAG;

*Fzd5*, CTGTGGTCTGTGCTGTGCTT and GGCCATGCCAAAGAAATAGA;

*Fzd6*, TCTGTGCCTCTGCGTATTTG and TCTCCCAGGTGATCCTGTTC;

*Fzd7*, GCTTCCTAGGTGAGCGTGAC and AACCCGACAGGAAGATGATG;

*Fzd8*, TTACATGCCCAACCAGTTCA and CGGTTGTAGTCCATGCACAG;

*Fzd9*, AGTTTCCTCCTGACCGGTTT and TTTTCGGTAGCACAGGCTCT;

*Fzd10*, AGATTCCCATGTGCAAGGAC and AGTTGGGGTCGTTCTTGTTG;

*Ror1*, GTGAAGTGCTGGAGAATGTC and GTCTACACCCGTGCTATTGT;

*Ror2*, CCACTGGGGTTCTATATGTG and CTGTGAACACTGGTCTGACA;

*Il1b*, GATCCCAAGCAATACCCAAAGAAG and CTCTGCTTGTGAGGTGCTGATGTA;

*Il6*, TAGTCCTTCCTACCCCAATTTCCA and ATGAATTGGATGGTCTTGGTCCTT;

*Il10*, AAGCCTTATCGGAAATGATCCAGT and CTTCTCACCCAGGGAATTCAAATG;

*Tnfa*, CTGGGACAGTGACCTGGACTGT and ACTCTCCCTTTGCAGAACTCAGG;

*Ifng*, CTTTAACAGCAGGCCAGACA and GCGAGTTATTTGTCATTCGG;

*Mmp1a*, CCCATATGCCATTACTCACAACAA and AACTGCTTTTGGCAAATATGGTGT;

*Mmp1b*, CAGATCCTGAAACCCTGAGAGCTA and GTCCAACGAGGATTGTTGTGAGTA;

*Mmp3*, AGAAGATCGATGCTGCCATTTCTA and TCCATGGATTGTTTCTTCTCATCA;

*Gdf15*, CTTGAAGACTTGGGCTGGAG and TAAGAACCACCGGGGTGTAG;

*Edn3*, GCAGGTCTGGGAAACAAGAG and CTGGGAGCTTTCTGGAACTG;

*Tgfb2*, CAGCGCTACATCGATAGCAA and CCTCGAGCTCTTCGCTTTTA;

*Tbp*, GAATAAGAGAGCCACGGACAACTG and AAGCCCAACTTCTGCACAACTCTA.

**Flow cytometry**

Hearts were excised from mice and perfused with sterile PBS to remove residual blood. The tissues were minced into ~1 mm pieces and digested at 37 °C in 5 ml of digestion buffer (833.33 µl of 4 mg/ml Liberase TH, 300 µl of 1 M HEPES, 30 µl of 10 mg/ml DNAse, 28.84 ml of Hanks Balanced Salt Solution (HBSS)). After 10 minutes, the supernatant was filtered through a 100 µm cell strainer and fresh digestion buffer was added. This procedure was repeated three times, followed by flushing with digestion buffer.

Cardiac cells were collected, and cardiomyocytes were separated from non-myocytes by gravity sedimentation for 20 minutes at room temperature. Non-myocyte pellets were recovered by centrifugation and washed with MACS buffer (Sterile PBS supplemented with 0.5% BSA and 1 mM EDTA). Single-cell suspensions were resuspended in 1X RBC lysis buffer (420302, BioLegend) and filtered through a 40 µm cell strainer. To minimize non-specific binding, cells were incubated with anti-CD16/32 FcReceptor (FcR) blocking antibody (101330, BioLegend), followed by staining with the Zombie Agua Fixable Viability Kit (42310, BioLegend). Cells were then incubated with fluorescent-conjugated antibodies against cell surface antigens or appropriate isotype controls (antibody details are provided below). Flow cytometry and cell sorting were performed on a FACSAria II instrument using BD FACSDiva software v8.0.2 (BD Biosciences).

The following antibodies were used for staining: APC Mouse IgG1, κ Isotype Ctrl (400119, BioLegend); PerCP/Cyanine5.5 Rat IgG2a kappa Isotype Ctrl Antibody, clone RTK2758 (400531, BioLegend); PE Mouse IgG1, κ Isotype Ctrl (12-4714-81, Invitrogen); APC Mouse feeder cells antibody, clone mEF-SK4 (130-120-802, Miltenyi Biotech); PerCP-eFluor 710 Mouse CD31 (PECAM-1) (46-0311-80, Invitrogen); PE Mouse CD45, clone 30-F11 (103106, BioLegend).

**Immunoblotting**

Mouse heart homogenates and cell lysates were prepared in RIPA lysis buffer (89900, Thermo Fisher Scientific) supplemented with protease and phosphatase inhibitors (78425, Sigma-Aldrich). Protein concentrations were determined using the Pierce™ BCA Protein Assay Kit (23225, Thermo Fisher Scientific) according to the manufacturer's instructions. Equal amounts of protein were separated by SDS-PAGE and transferred to PVDF membranes (10120, VWR). Membranes were incubated with the indicated primary antibodies (see Antibodies and reagents section), and immunoreactive bands were detected using an enhanced chemiluminescence substrate (ECL Prime Western Blotting Detection Reagent, GE Healthcare) and visualized with a Bio-Rad ChemiDoc imaging system. Band intensities were quantified by densitometry using ImageJ software.

**Antibodies and reagents**

Wnt5a (1:1000, MA5-14946, Thermo Fisher Scientific), p21 (1:500, sc-6246, Santa Cruz Biotechnology), p53 (1:500, sc-126, Santa Cruz Biotechnology), PML (H-238) (1:500, sc-5621, Santa Cruz Biotechnology), Phospho-Histone H2A.X (Ser139) (20E3) (1:1000, 9718S, Cell Signaling), αTubulin (1:3000, 2144S, Cell Signaling), cTnT (1:2000, MA5-12960, Thermo Fisher Scientific), Phospho-p53 (Ser15) (1:1000, 9284S, Cell Signaling), Vinculin (1:3000, V9131, Sigma-Aldrich), Gapdh (14C10) (1:3000, 2118L, Cell Signaling), Fzd2 (1:1000, 24272-1-AP, Proteintech).

The siRNAs used in this study are as follows: rat *Wnt5a* siRNA (identifier (ID): s134311), rat *Fzd2* siRNA (ID: s134162), rat *Pml* siRNA (ID: 289378), siNegative Control (4390843) (Silencer Select pre-designed siRNA from Thermo Fisher Scientific) and Non-targeting control pool (D-001810-10-05) (ON-TARGET plus SMART pool from Horizon Discovery Biosciences (Dharmacon)). The transfection reagent used is lipofectamine RNAiMAX (cat no. 13778075, Invitrogen) or DharmaFECT 1 Transfection Reagent (T-2001-01, Horizon Discovery Biosciences (Dharmacon)).

Recombinant human/mouse Wnt5a (645-WN) and recombinant human sFRP5 (6266-SF-050) were purchased from R&D Systems.

**Supplementary Data**

**Table S1. Clinical characteristics of patients with paired serum samples collected before and approximately 3 months after anthracycline therapy.**

| Patient ID | Age (years) | Sex | Diagnosis | Anthracycline regimen |
|---|---|---|---|---|
| VV1 | 47 | Female | Breast cancer | Epirubicin<br>151.2 mg/$m^2$ x 4 cycles |
| VV2 | 43 | Female | Breast cancer | Epirubicin<br>140 mg/$m^2$ x 3 cycles |
| VV5 | 39 | Female | Breast cancer | Epirubicin<br>159.3 mg/$m^2$ x 4 cycles |
| VV11 | 64 | Female | Breast cancer | Epirubicin<br>135 mg/$m^2$ x 4 cycles |
| VV13 | 49 | Female | Breast cancer | Epirubicin<br>154.80 mg/$m^2$ x 4 cycles |
| VV15 | 59 | Female | Breast cancer | Doxorubicin<br>96 mg/$m^2$ x 4 cycles |
| VV16 | 51 | Female | Breast cancer | Epirubicin<br>162.9 mg/$m^2$ x 4 cycles |
| VV17 | 42 | Female | Breast cancer | Epirubicin<br>176.4 mg/$m^2$ x 4 cycles |
| VV22 | 73 | Female | Breast cancer | Epirubicin<br>144 mg/$m^2$ x 4 cycles |
| VV23 | 56 | Female | Breast cancer | Doxorubicin + paclitaxel<br>105 mg/$m^2$ x 4 cycles |
| VV25 | 53 | Female | Breast cancer | Epirubicin + paclitaxel<br>150 mg/$m^2$ x 4 cycles |
| VV27 | 47 | Female | Breast cancer | Epirubicin<br>163.80 mg/$m^2$ x 4 cycles |
| VV28 | 74 | Female | Breast cancer | Epirubicin<br>160.20 mg/$m^2$ x 4 cycles |
| VV30 | 61 | Female | Breast cancer | Epirubicin<br>148.5 mg/ $m^2$ x 4 cycles |
| VV35 | 64 | Female | Breast cancer | Epirubicin + paclitaxel<br>171.90 mg/ $m^2$ x 4 cycles |
| VV36 | 66 | Female | Breast cancer | Doxorubicin<br>91 mg/$m^2$ x 4 cycles |
| VV38 | 56 | Female | Breast cancer | Epirubicin<br>162 mg/$m^2$ x 4 cycles |

Paclitaxel was administered as part of the chemotherapy regimen where indicated.

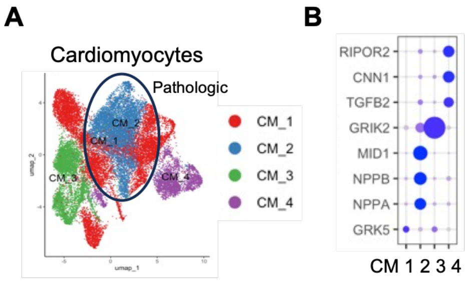


**Figure S1. Cardiomyocyte-state annotation in human DOX-induced cardiomyopathy snRNA-seq dataset.** (**A**) UMAP visualization of cardiomyocyte subclusters from donor and DOX-induced cardiomyopathy hearts in a published human snRNA-seq dataset. Cardiomyocyte states were annotated as CM_1, CM_2, CM_3 and CM_4. The pathologic cardiomyocyte population is indicated. (**B**) Dot plot showing representative marker genes used to annotate cardiomyocyte states. Dot size indicates the fraction of cells expressing each gene and color indicates mean expression.

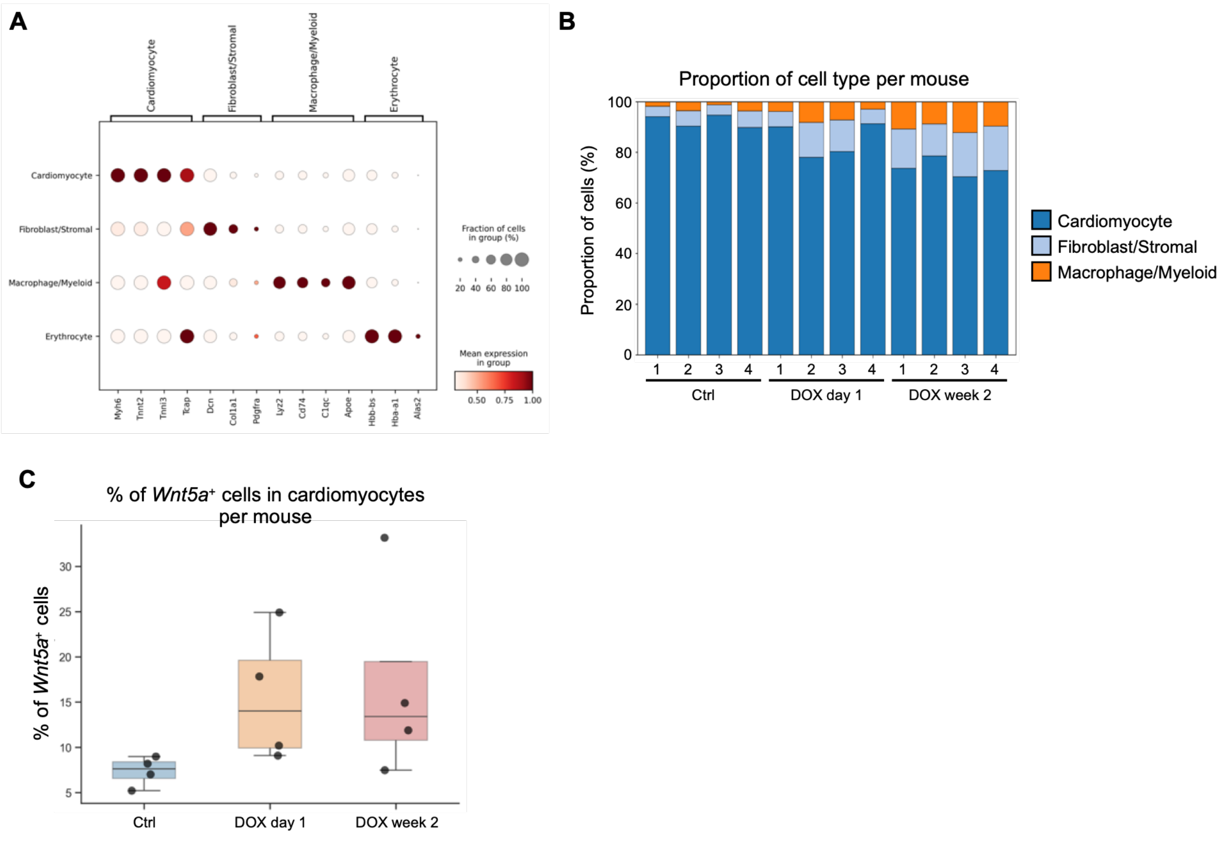


**Figure S2. Mouse heart scRNA-seq annotation and *Wnt5a* expression in cardiomyocytes after DOX treatment.** (**A**) Dot plot showing representative marker genes used to annotate major cardiac cell populations in mouse heart scRNA-seq analysis. (**B**) Proportion of major cardiac cell types per mouse in control, DOX day 1 and DOX week 2 hearts. (**C**) Mouse-level quantification of the percentage of Wnt5a-positive cardiomyocytes in control, DOX day 1 and DOX week 2 hearts. Each dot represents one mouse.

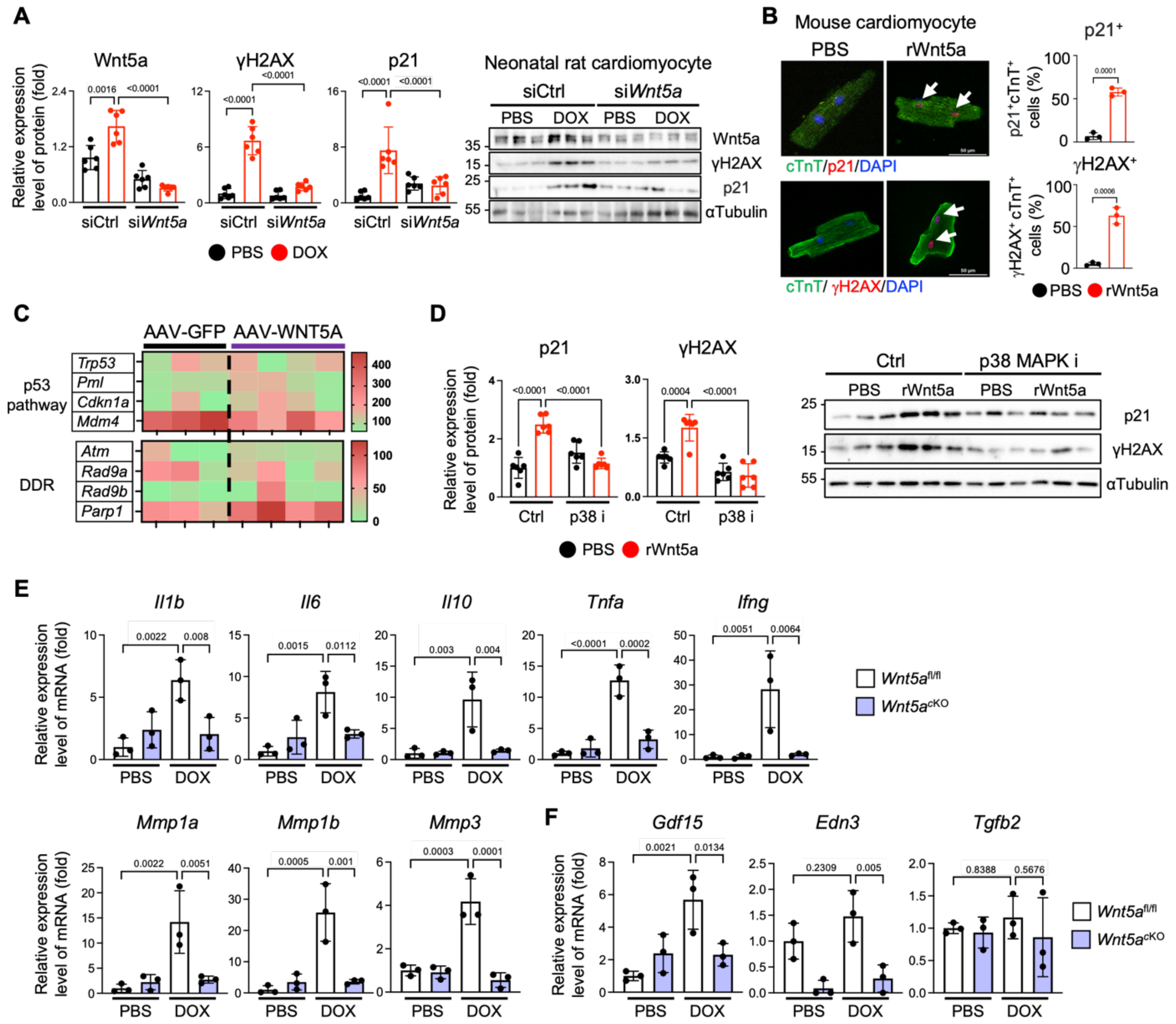


**Figure S3. Wnt5a promotes senescence-associated signaling and SASP-related gene expression in cardiomyocytes.** (**A**) NRVMs were transfected with control siRNA or *Wnt5a* siRNA and treated with PBS or DOX (100 nM for 48 hours). Cell lysates were subjected to Western blotting with anti-Wnt5a, γH2AX, p21 and αTubulin antibodies. Relative expression levels of proteins were normalized to αTubulin. (**B**) Representative immunofluorescence images and quantification of p21-positive and γH2AX-positive adult mouse cardiomyocytes treated with PBS or rWnt5a (50 ng/ml for 48 hours). Cells were stained for cTnT, p21 or γH2AX and with DAPI. Arrows indicate p21-positive or γH2AX-positive nuclei in cTnT-positive cardiomyocytes. Scale bars: 50 μm. (**C**) Heatmap from bulk RNA-seq showing expression of p53 pathway and DNA damage response genes in hearts from mice injected with AAV9-*cTNT-GFP* or AAV9-*cTNT-WNT5A*. (**D**) NRVMs were treated with PBS or rWnt5a (50 ng/ml) in the presence or absence of a p38 MAPK inhibitor (5 μM) for 48 hours. Cell lysates were subjected to Western blotting with γH2AX, p21 and αTubulin antibodies. Relative expression levels of proteins were normalized to αTubulin. (**E**) qPCR analysis of conventional SASP-related genes and matrix-remodeling SASP genes in hearts from *Wnt5a*$^{fl/fl}$ and *Wnt5a*$^{cKO}$ mice treated with PBS or DOX. (**F**) qPCR analysis of atypical SASP-related genes in hearts from *Wnt5a*$^{fl/fl}$ and *Wnt5a*$^{cKO}$ mice treated with PBS or DOX. n = 3. Data are presented as mean ± SD. Statistical significance was determined by two-way ANOVA followed by Sidak's multiple-comparison test (**A**, **D**, **E**, **F**) and by unpaired Student's t-test (**B**). Exact P values are shown in the graphs.

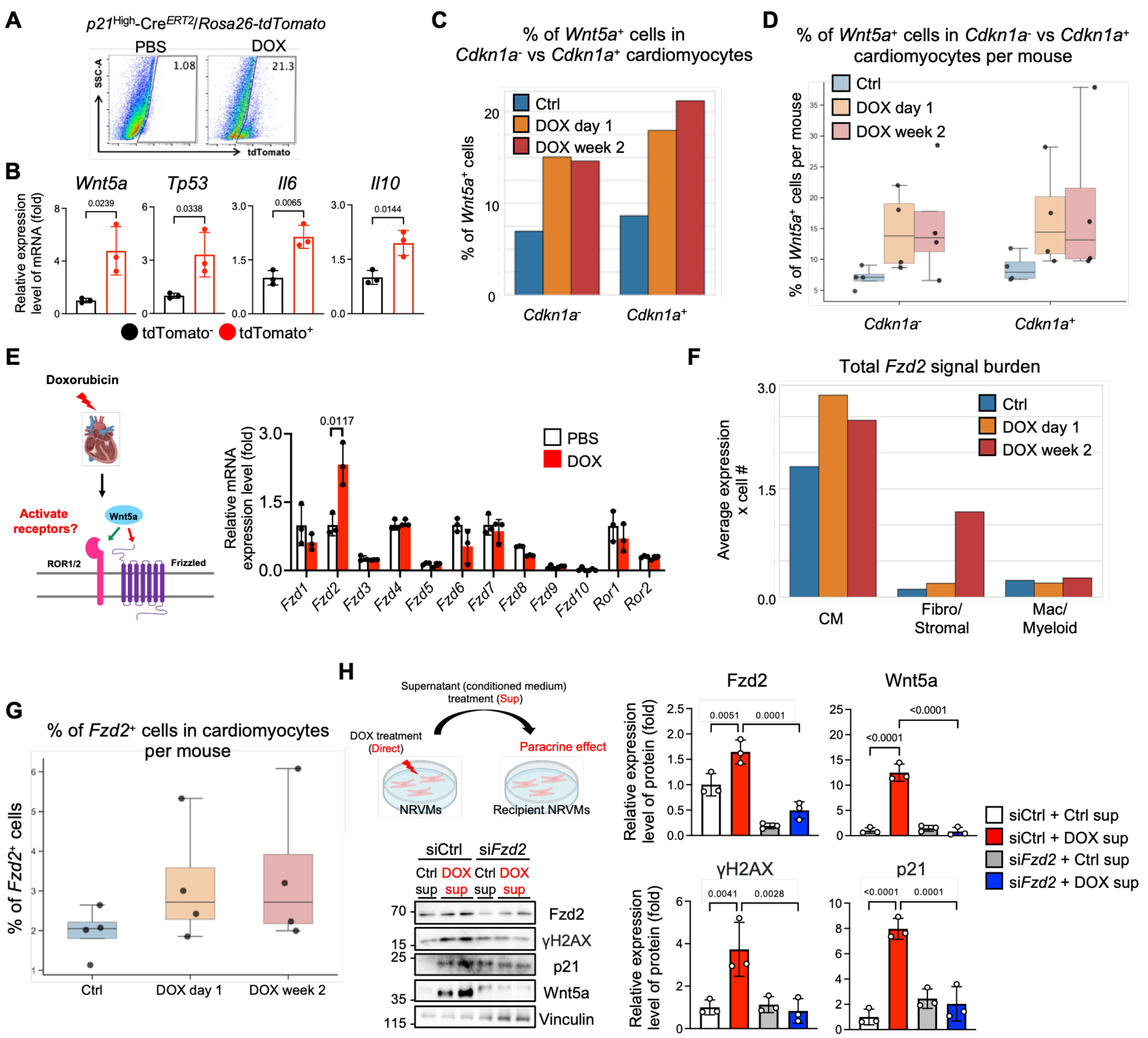


**Figure S4. *Wnt5a* expression in *Cdkn1a*-positive cardiomyocytes and Fzd2-dependent paracrine signaling after DOX treatment.** (**A**) Representative flow cytometry plots showing tdTomato-positive cardiomyocytes from *p21*$^{High}$-Cre$^{ERT2}$;*Rosa26-tdTomato* mice treated with PBS or DOX (cumulative 20 mg/kg). (**B**) tdTomato-negative or tdTomato-positive cardiomyocytes were sorted from DOX-treated *p21*$^{High}$-Cre$^{ERT2}$;*Rosa26-tdTomato* mice and subjected to qPCR analysis. (**C**) Pooled cell-level quantification from mouse heart scRNA-seq data showing the percentage of *Wnt5a*-positive cells within *Cdkn1a*-negative and *Cdkn1a*-positive cardiomyocyte populations in control, DOX day 1, and DOX week 2 hearts. (**D**) Mouse-level quantification from mouse heart scRNA-seq data showing the percentage of *Wnt5a*-positive cells within *Cdkn1a*-negative and *Cdkn1a*-positive cardiomyocyte populations in control, DOX day 1, and DOX week 2 hearts. Each dot represents one mouse. (**E**) qPCR analysis of Wnt receptor expression in mouse hearts treated with PBS or DOX. (**F**) Quantification of the total *Fzd2* signal burden across major cardiac cell populations in the mouse heart scRNA-seq analysis, calculated as average *Fzd2* expression × number of *Fzd2*-expressing cells. (**G**) Mouse-level quantification from mouse heart scRNA-seq data showing the percentage of *Fzd2*-positive cardiomyocytes in control, DOX day 1, and DOX week 2 hearts. Each dot represents one mouse. (**H**) NRVMs were transfected with control siRNA or *Fzd2* siRNA and treated with conditioned medium (sup) from control (PBS) or DOX-treated cardiomyocytes for 48 hours. Schematic of conditioned medium transfer experiments and immunoblot analyses with quantification of Fzd2, Wnt5a, γH2AX, and p21 protein levels in recipient cardiomyocytes. Relative expression levels of proteins were normalized to Vinculin. Data are presented as mean ± SD. Statistical significance was determined by unpaired Student's t-test (**B**, **E**) and by one-way ANOVA followed by Tukey's multiple-comparison test (**H**). Exact P values are shown in the graphs.

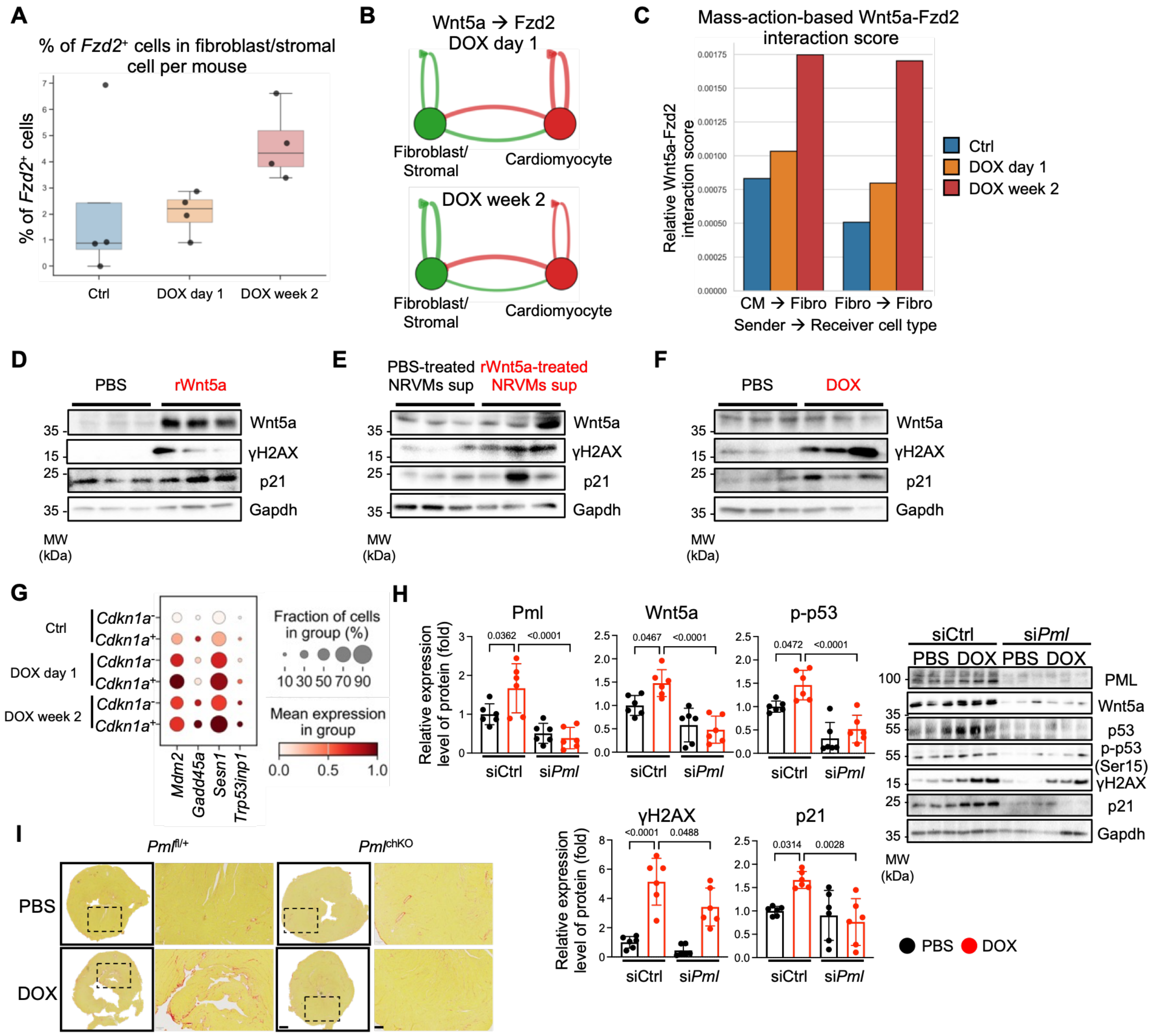


**Figure S5. Wnt5a-Fzd2 signaling promotes cardiomyocyte-fibroblast communication and Pml-dependent senescence signaling.** (**A**) Mouse-level quantification from mouse heart scRNA-seq data showing the percentage of *Fzd2*-positive fibroblast/stromal cells in control, DOX day 1, and DOX week 2 hearts. Each dot represents one mouse. (**B**) Inferred *Wnt5a-Fzd2* communication map between cardiomyocytes and fibroblast/stromal cells at DOX day 1 and DOX week 2. Arrow thickness represents relative interaction strength. (**C**) Mass-action–based *Wnt5a-Fzd2* interaction scores for cardiomyocyte-to-fibroblast/stromal and fibroblast/stromal-to-fibroblast/stromal signaling in control, DOX day 1, and DOX week 2 hearts. (**D**) Neonatal rat cardiac fibroblasts (NRCFs) were treated with PBS or rWnt5a (50 ng/ml) for 48 hours. Cell lysates were subjected to Western blotting with anti-Wnt5a, γH2AX, p21 and Gapdh antibodies. (**E**) NRCFs were treated with conditioned medium from PBS- or rWnt5a (50 ng/ml for 48 hours)-treated NRVMs. (**F**) NRCFs were treated with PBS or DOX (100 nM) for 48 hours. Cell lysates were subjected to Western blotting with anti-Wnt5a, γH2AX, p21 and Gapdh antibodies. (**G**) Dot plot from mouse heart scRNA-seq data showing expression of selected p53 target and stress-response genes in *Cdkn1a*-negative and *Cdkn1a*-positive cardiomyocytes from control, DOX day 1, and DOX week 2 hearts. Dot size indicates the fraction of cells expressing each gene, and color indicates mean expression. (**H**) NRVMs were transfected with control siRNA or si*Pm* and treated with PBS or DOX (100 nM) for 48 hours. Cell lysates were subjected to Western blotting with anti-PML, Wnt5a, p53, phospho-p53, γH2AX, p21 and Gapdh antibodies. Protein levels were normalized by Gapdh. (**I**) Representative Picric Sirius Red–stained heart sections from $Pml^{fl/+}$ and $Pml^{chKO}$ mice treated with PBS or DOX (cumulative 20 mg/kg). Data are presented as mean ± SD. Statistical significance was determined by two-way ANOVA followed by Sidak's multiple-comparison test (**H**). Exact P values are shown in the graphs.

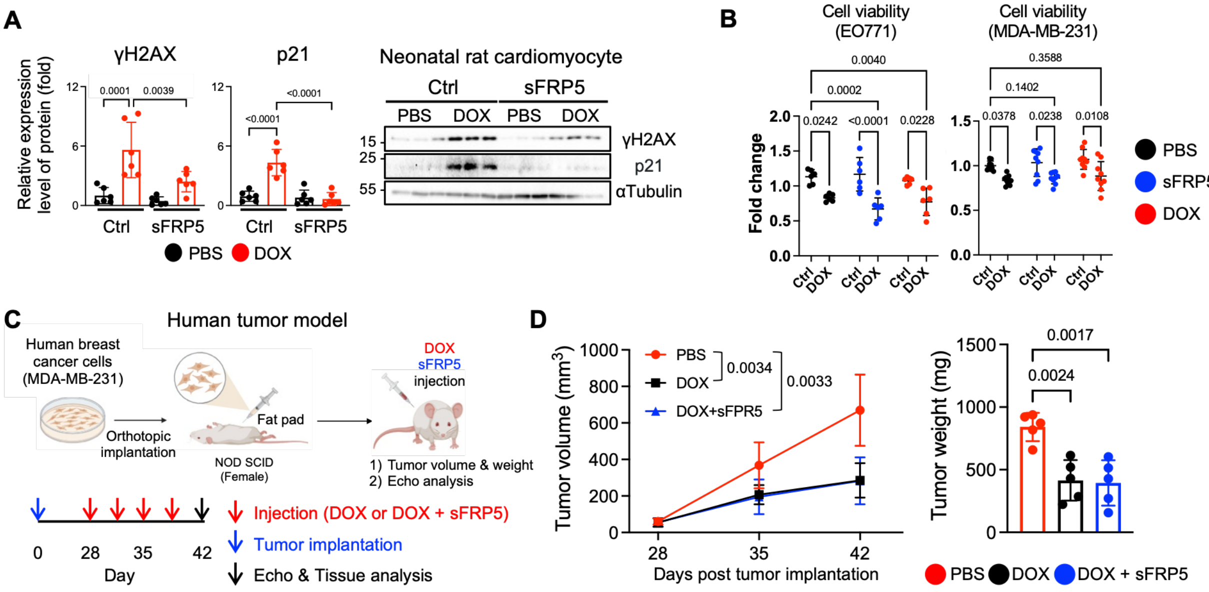


**Figure S6. sFRP5 suppresses DOX-induced cardiomyocyte senescence-associated markers and preserves DOX anti-tumor activity in human breast cancer models.** (**A**) NRVMs were treated with PBS or DOX (100 nM) in the presence or absence of recombinant sFRP5 (400 ng/ml) for 48 hours. Cell lysates were subjected to Western blotting with anti-γH2AX, p21 and αTubulin antibodies. Relative expression levels of proteins were normalized to αTubulin. (**B**) Mouse EO771 and human MDA-MB-231 breast cancer cells were treated with PBS or DOX (500 nM) in the presence or absence of recombinant sFRP5 (400 ng/ml) or WNT5A (100 ng/ml). Cell viability analysis was performed after 48 hours treatment. (**C**) Schematic of the human MDA-MB-231 orthotopic breast cancer model and treatment protocol. MDA-MB-231 cells were implanted into the mammary fat pads of female NOD SCID mice, followed by treatment with PBS, DOX (5 mg/kg, i.p.), or DOX plus sFRP5 (20 μg/kg, i.p.). Tumor growth, tumor weight, cardiac function, and tissue analysis were assessed at the indicated time points. (**D**) Tumor volume over time and final tumor weight in MDA-MB-231 tumor-bearing mice treated with PBS, DOX, or DOX plus sFRP5. Data are presented as mean ± SD. Statistical significance was determined by two-way ANOVA followed by Sidak's multiple-comparison test (**A** and **B**), and by one-way ANOVA followed by Tukey's multiple-comparison test for endpoint tumor measurements (**D**). For **D**, statistical comparisons were performed using tumor volumes at the final measurement time point. Exact P values are shown in the graphs.
